# Transcriptional competition biases the effects of second messengers in *Escherichia coli*

**DOI:** 10.1101/2025.04.15.648722

**Authors:** Andrea Ripamonti, Milan Lacassin, Rossana Droghetti, Gregory Bokinsky, Marco Cosentino Lagomarsino

**Author notes:** These authors contributed equally (Andrea Ripamonti, Milan Lacassin). These authors contributed equally (Gregory Bokinsky, Marco Cosentino Lagomarsino).

## Abstract

Cells regulate gene expression by balancing transcriptional resources across different functional groups of genes. In *Escherichia coli*, second messengers such as ppGpp and cAMP control ribosome biogenesis and metabolic gene expression, respectively. While these regulators are typically studied in isolation, we provide a theory showing that their effects are intertwined due to global transcriptional competition. Using experimental data from RelA overexpression and a mechanistic modeling framework, we show that ppGpp-mediated repression of ribosomal genes competes for transcriptional resources with cAMP-driven activation of catabolic genes. This competition reshapes proteome allocation in a way that transcends individual regulators. Our findings challenge common modeling assumptions about transcription factor action and revive classical ideas suggesting that large-scale gene regulation should be studied within the broader context of resource availability, with implications for understanding cellular regulation across diverse biological systems, beyond bacterial physiology.

## INTRODUCTION

Precise regulation of gene expression is fundamental to cellular physiology, with implications for fitness and disease^1–4^. For instance, cells respond to growth limitations by expressing ribosomes and enzymes required to balance the supply and demand of key metabolites^5^. In this context, the proteome sector theory has emerged, particularly in *Escherichia coli* ^6–10^, as a powerful mathematical approach that captures large-scale gene regulatory principles due to mass balance and supply-demand, including the well-established linear relationship between ribosomal content and growth rate (sometimes termed the “ribosomal growth law”^6,11^). Recent advances in proteomics have further refined our understanding of proteome allocation, revealing how cells can prioritize protein synthesis in response to environmental conditions and internal state^8,9,12–15^.

A crucial open question is how cells integrate environmental signals and internal sensing to regulate ribosomal allocation and change the growth rate. One key mechanism for growth rate control in *E. coli* involves the alarmone (p)ppGpp (henceforth shortened as ppGpp), a global regulator of the ribosomal protein sector^1^. Together with the transcription factor DksA, ppGpp binds to RNA polymerase (RNAP) inducing a conformational change that inhibits the transcription of genes encoding ribosomal RNA and proteins^16–19^. Additionally, ppGpp has many intracellular targets potentially linked to metabolism and translation, but the transcriptional effect appears to be dominant for ribosome regulation^16^.

Recent work by Wu et al.^20^ demonstrated that ppGpp levels quantitatively reflect translation elongation rates, establishing a direct link between the cellular perception of growth rate and ribosomal biogenesis. Mechanistically, ppGpp is synthesized from GTP by the ribosome-bound enzyme RelA when uncharged tRNAs bind to the ribosomal A site^21–23^. The enzyme SpoT also has a limited capacity for ppGpp synthesis, though its main role is as a ppGpp hydrolase^24^. Thus, under starvation, ppGpp acts as a sensor of nutrient availability and redirects resources from the translation machinery towards metabolism^25,26^.

Just as cellular resources must be efficiently allocated at the protein level, transcription itself requires careful distribution of RNA polymerase across different genes. The limited transcriptional capacity of the RNAP pool represents a fundamental constraint on gene expression, analogous to the limited proteome capacity^27–29^. This constraint means that increased transcription of one gene or set of genes necessarily affects the expression of others through competition for RNAP. While traditional views of transcriptional regulation focus on individual promoter-regulator interactions, the allocation of RNAP across different promoters emerges as a critical systems-level consideration, particularly when examining large-scale changes in gene expression such as those involving entire proteome sectors^30,31^.

This perspective suggests that global regulators like ppGpp might coordinate cellular responses not only through direct promoter interactions but also by influencing the competition for RNAP allocation across the genome. Thus, their effect should depend on the contextual activity of other global regulators. One such example is the second messenger cyclic AMP (cAMP), a small molecule that modulates metabolic gene expression in response to the availability of carbon substrates^32,33^. Despite their distinct roles, both ppGpp and cAMP influence proteome allocation at the transcriptional level, shaping the ribosomal and metabolic sectors in response to nutrient availability. While specific crosstalk between (p)ppGpp and other nucleotide messengers is being explored^34^, including signal conversion, allosteric regulation, and target competition, competition for RNAP allocation has comparatively received less attention.

Perturbing these second messengers by “opening the loops” of feedback control^35,36^ provides a powerful strategy to address this gap, allowing us to test our understanding of transcriptional regulation and gain key insights into regulatory interactions. For instance, Zhu and Dai^37^ explored the effects of ppGpp overabundance by overexpressing its synthase enzyme, RelA, demonstrating that the ribosomal growth law is robust, despite broad proteomic rearrangements^38,39^. Similarly, Kochanowski et al.^40^ showed that cAMP titration preserves the same growth law. While these findings highlight the robustness of the ribosomal growth law, they also raise fundamental questions about the mechanisms governing proteome allocation under these perturbations. Intriguingly, prior studies suggest that the link between ppGpp and the growth rate is not univocal, particularly when RelA is overexpressed^41–43^. This divergence (reviewed in ref.^44^) has so far remained unexplained and challenges the prevailing framework, suggesting a more complex interplay between ppGpp, global transcriptional regulation, and growth rate control.

Although recent models offer promising steps toward a mechanistic theory of proteome allocation, key gaps remain. Existing theories that connect proteome sectors with nutrient and growth sensing often overlook the mechanistic details of transcription^36,45,46^. Recent studies by Balakrishnan et al. and by Lin and Amir^27,31^ provide a solid foundation for describing the flow of information from DNA to proteins, yet they are not fully embedded within the proteome sector framework. Conversely, the model proposed by Calabrese et al.^29^ incorporates transcription and translation but lacks the regulatory details needed to capture the effects of second messengers.

In this work, we address the limitations of existing theories by adopting a theory-first approach. Building on the study of Balakrishnan et al., we develop a framework that integrates transcriptional regulation by second messengers and competition for RNA polymerases into proteome sector models. This framework enables a model-guided analysis of experimental data. Leveraging published datasets from refs.^20,31,37–39,47^, as well as performing new experiments involving RelA overexpression, we investigate a previously uncharacterized divergence in the relationship between ppGpp and ribosomal content. By simultaneously monitoring the levels of ppGpp and cAMP, we uncover non-trivial effects of second messengers on proteome coordination. These effects are effectively captured by our model, which describes second messen-ger–regulated transcriptional competition between proteome sectors.

## RESULTS

### Global transcriptional coupling connects differentially regulated proteome sectors

Building on the framework of Balakrishnan et al.^31^, we introduce a model that integrates transcriptional regulation by second messengers and competition for RNA polymerases (RNAPs) into proteome sector dynamics (see Methods S1 for detailed derivations and assumptions). We denote the mass fraction of each proteome sector *I* as

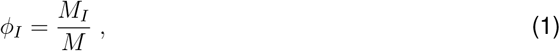

where *M*_*I*_ is the total mass of proteins in sector *I* and *M* is the total protein mass. For example, the “ribosomal sector” (*ϕ*_*R*_) includes proteins involved in translation, while other sectors, such as metabolic (*ϕ*_*P*_) and housekeeping (*ϕ*_*Q*_), are categorized based on their response to growth limitations^6,8,9,33^.

The model focuses on how mRNAs from different sectors are produced. We define the “transcriptome fraction” *χ*_*I*_ as the fractional proportion of mRNAs encoding proteins in sector *I*. Assuming no post-transcriptional regulation, the time evolution of protein mass fractions is given by (Methods S1 for derivation)

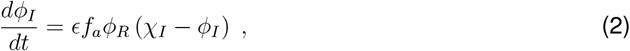

where *ϵ* is the translation elongation rate, *f*_*a*_ is the fraction of active ribosomes, and the product *ϵf*_*a*_*ϕ*_*R*_ corresponds to the growth rate *λ*.

Transcript dynamics are described in an “initiation-limited” regime^31,48^, where the concentration of a transcript species [*m*_*i*_] evolves as

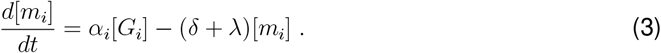

Here *α*_*i*_ is the transcription initiation rate, *δ* is the uniform mRNA degradation rate and [*G*_*i*_] is the gene concentration. The dilution term − *λ*[*m*_*i*_] accounts for the time variation in cellular volume. Initiation rates depend on the concentration of available RNAP holoenzymes [*N*_*av*_], and are given by

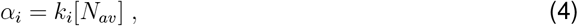

where *k*_*i*_ is the promoter strength, influenced by transcriptional regulators.

Balakrishnan et al.^31^ found that holoenzyme formation is limited by sigma factors, mainly by the housekeeping RpoD (*σ*^70^), rather than by RNAP core components. Therefore, as a first approximation, the above equation can be interpreted as reflecting the assumption of *σ*^70^-limited transcription initiation. Although the model can be extended to incorporate alternative sigma factors (Methods S1), we do not expect them to significantly affect our main results (see Discussion). Similarly, while core RNAP abundance can be described with a dedicated proteome sector^27–29^, our simplified framework does not rely on it. We emphasize that a non-RNAP-limited regime is possible^49^, but it falls outside the scope of this study, since in this regime transcription output per gene is constant and total mRNA scales directly with the number of promoters^27^.

The key novelty of our approach lies in embedding transcription into proteome sector theory by introducing sector-specific coarse-grained promoter strengths *β*_*I*_. This allows us to express mRNA dynamics at the sector level

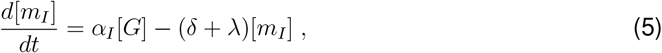

where we defined the “coarse-grained initiation rate” *α*_*I*_ = *β*_*I*_[*N*_*av*_] and the typical gene concentration [*G*], which we assume to be uniform across all sectors (see STAR Methods and Methods S1 for a model variant including gene dosage effects). From this, we derive the time evolution of transcriptome fractions *χ*_*I*_ (Methods S1)

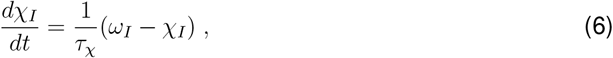

where 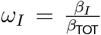 represents RNAP allocation across sectors, with *β*_TOT_ = ∑_*J*_ *β*_*J*_, while *τ*_*χ*_ is a characteristic relaxation timescale.

In steady-state growth, our model predicts that proteome fractions are determined transcriptionally by sector “strengths” as follows,

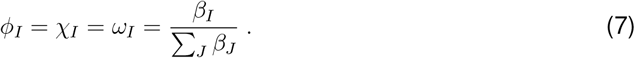

This relationship, proposed for single genes in ref.^31^, implies that transcriptional regulation in one sector indirectly affects other sectors due to the normalization condition ∑_*I*_ *ω*_*I*_ = 1 (Fig. 1D). This coupling is central to our framework, since it sets how global transcriptional regulation and competition for RNAPs shape proteome coordination.

**Figure 1.**
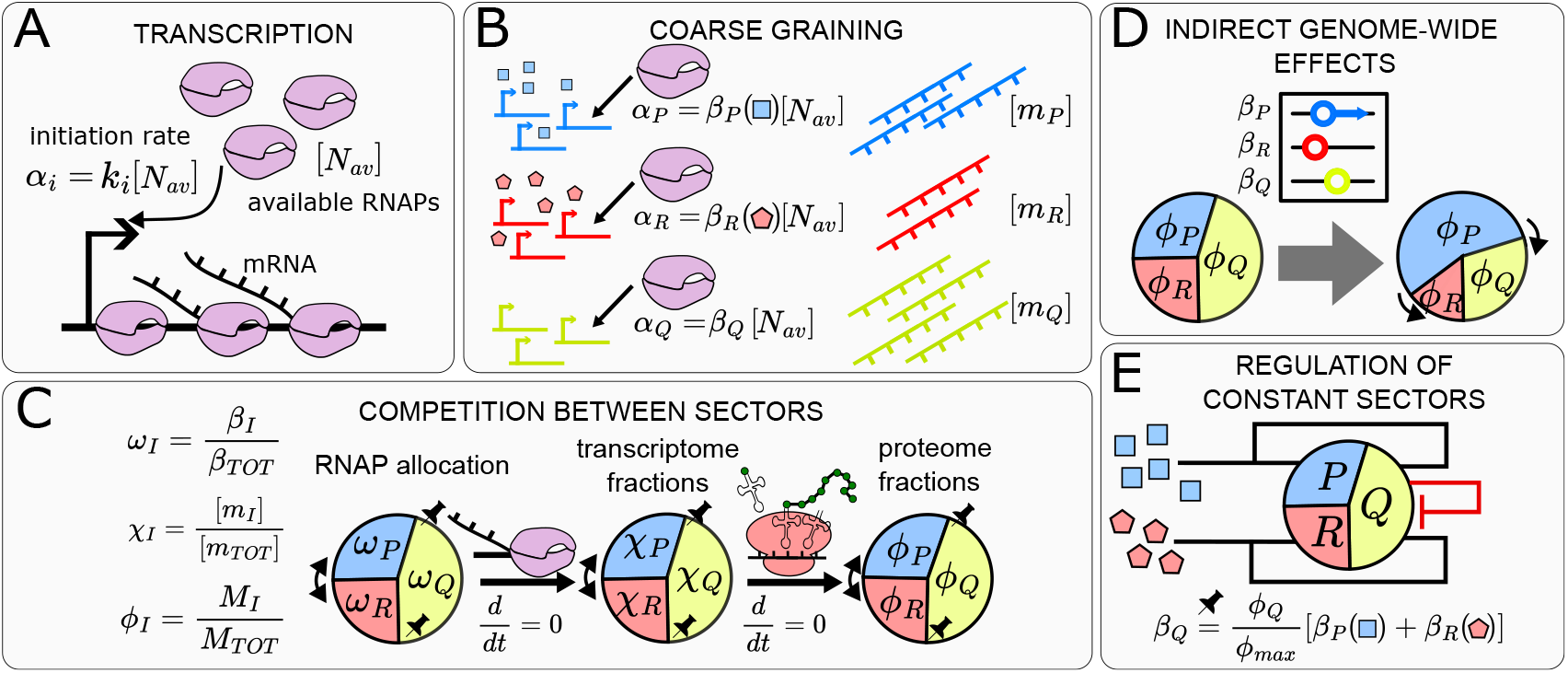
Illustration of the model for transcriptional competition and its features. (A) Transcription initiation rates are proportional to the concentration of available RNA polymerases [*N*^*av*^] (the model neglects traffic of RNA polymerase on genes). (B) Genes are grouped into sectors (three in this illustration). The R and P sectors are controlled by regulators, which act on the “sector strengths” *β* ^*R*^ and *β*^*P*^ at the transcriptional level. mRNA abundances [*m*^*I*^] are determined by the coarse-grained initiation rates *α* ^*I*^. (C) Sector strengths determine the transcriptome fractions *χ* ^*I*^. The Q sector is assumed to be regulated in such a way that *ϕ* ^*Q*^ is “pinned” at a constant value across conditions. At steady-state, transcriptome fractions correspond to proteome fractions *ϕ* ^*I*^. (D) Due to the normalization constraint, activation or repression of a single sector indirectly affects all others, even if their sector strengths remain unchanged. (E) Maintaining a constant Q sector size requires direct regulation, influenced either by signals coupled to the R and P sectors (black arrows) or by negative autoregulation (red T-shaped arrow), ensuring homeostasis.

### Sectors with constant size must be actively maintained at a fixed expression level

To understand the role played by transcriptional competition in setting the sizes of different sectors, we focused on a commonly used three-sector model^6,7,36^. This model typically assumes that the mass fraction *ϕ*_*Q*_ of the housekeeping sector remains constant^6^. We incorporate this constraint into our framework using Eq. (7) finding

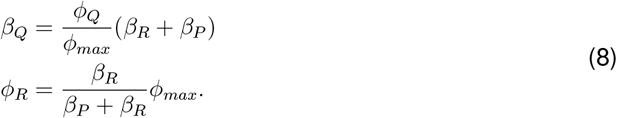

Since sectors P and R are regulated in order to adapt to different conditions (e.g. different nutrients), the sum *β*_*P*_ + *β*_*R*_ varies with the regulation, thus *β*_*Q*_ must follow its behaviour in a proportional way (Fig 1E). The second identity in Eq. (8) describes the competition between the P and R sectors for a proteome fraction equal to *ϕ*_*max*_:= 1 − *ϕ*_*Q*_. Note that eq. (8) strictly applies only at steady-state. In addition, the first relationship does not necessarily imply that the Q sector shares transcriptional regulators with the other two sectors. For instance, some studies^6,30,50^ have proposed that this sector maintains its constant size through negative autoregulation. Proteomics studies^8,9^ indicate that Q consists of multiple subsectors, each responding differently to growth limitations, while their overall sum remains constant. Regardless of the specific mechanisms governing Q, Eq. (8) provides the shape of its mean regulatory function in terms of *β*_*P*_ and *β*_*R*_.

### A positive covariation of ppGpp and cAMP coordinates proteome allocation under catabolic limitation

With a constant Q sector, proteome allocation is entirely determined by the relative size of R and P from equation (8), and ppGpp is believed to be a central driver of this process^16,21,51^ (Fig. 2A). To confirm this, we directly measured internal ppGpp levels in combination with growth rate across multiple conditions, corresponding to different carbon sources (see STAR Methods). We converted growth rate data into ribosomal fractions using the linear ribosomal growth law^6^. Consistent with the literature^20,37,41,52^, we observed a robust relationship between *ϕ*_*R*_ and *g*, with elevated ppGpp levels corresponding to slower growth rates, and thus smaller ribosomal fractions (Fig. 2B).

**Figure 2.**
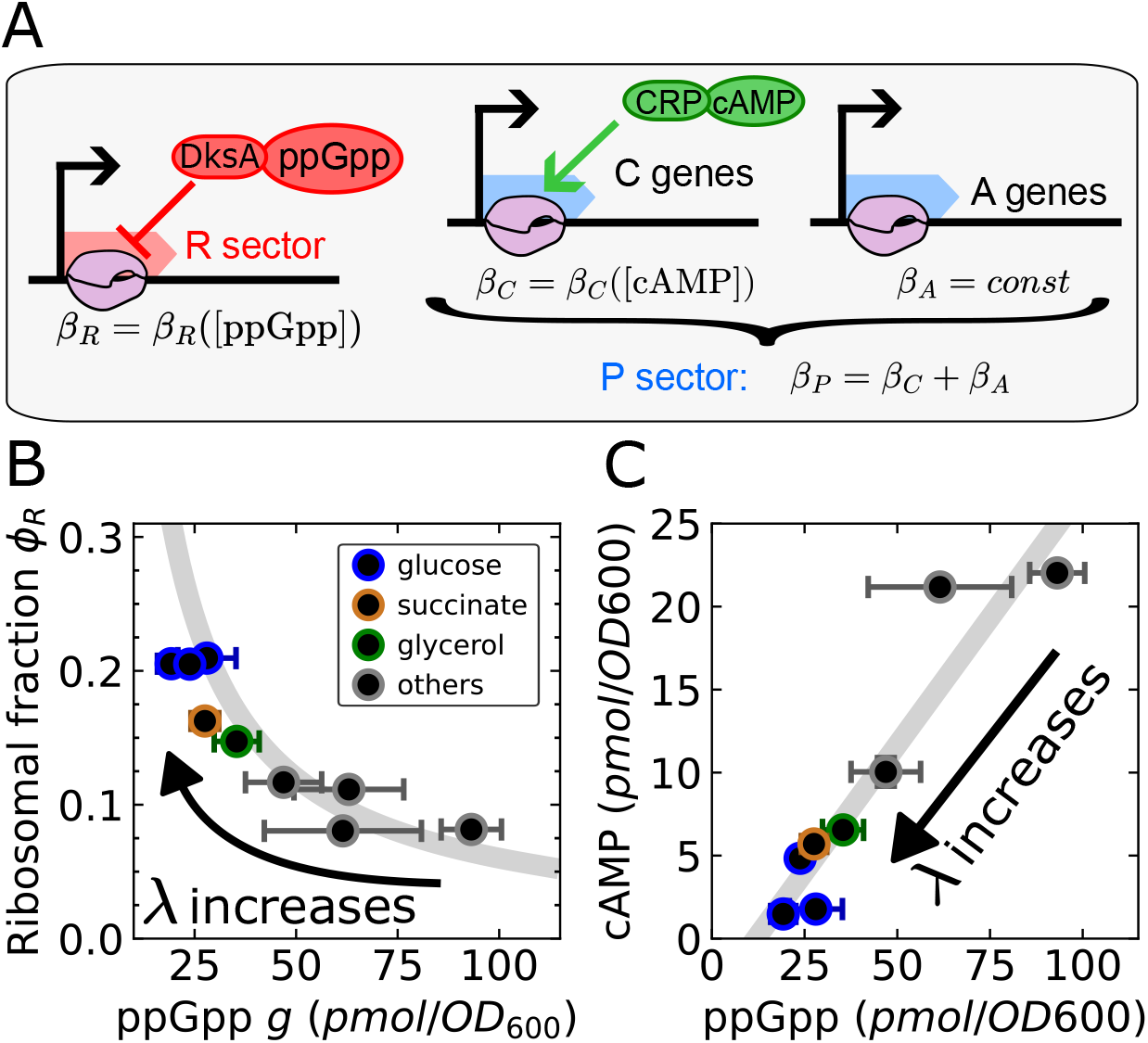
A positive covariation of ppGpp and cAMP under catabolic limitation defines proteome allocation as a function of ppGpp. (A) The model for transcriptional competition in *E. coli* describe the ribosomal (R) sector as repressed by ppGpp-DksA. This sector competes with the catabolic (C) sector, activated by cAMP-Crp, and with a constitutive (unregulated) anabolic (A) sector. C and A sectors form the P sector. (B) Under catabolic limitation, the ribosomal fraction *ϕ* ^*R*^ decreases with increasing ppGpp. The plot derives from our direct measurements of ppGpp levels and growth rate. Values for the ribosomal fraction are estimated from growth rate using joint measurements of RNA/protein ratio and growth rate from ref. ^37^. The gray line repre-sents the best fit according to the expression 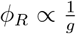 as in ref. ^20^. (C) Under catabolic limitation, cAMP increases approximately linearly with ppGpp (and therefore with decreasing growth rate) in the wild type (gray line, best fit). This correlation obscures the specific effect of cAMP on *ϕ*^*R*^ when considered solely as a function of ppGpp.

With only three proteome sectors, incorporating a passive P sector is sufficient to account for this growth rate-dependent proteome partitioning, consistently with the original suggestion by Scott et al.^6^. Under these assumptions, the steady-state proteome composition is a function of ppGpp only: *ϕ*_*R*_ = *ϕ*_*R*_(*g*) and *ϕ*_*P*_ = *ϕ*_*P*_ (*g*). Models with ppGpp as the sole transcriptional regulator are commonly found in the literature^20,36,45^, because this minimal setup effectively captures the correlation between *ϕ*_*R*_ and *g* across nutrient conditions seen in various *E. coli* studies.

However, previous studies^1,33,40,53–55^ show the importance of the second messenger cyclic AMP (cAMP) as a global regulator. cAMP activates the transcription factor CRP (cAMP receptor protein), which in turn drives catabolic gene expression by binding specific DNA sequences and interacting with the *α* subunit of RNA polymerase^32,56,57^. Produced by adenylate cyclase (Cya), cAMP levels depend on carbon substrate availability. You et al.^33^ proposed that Cya activity is repressed by alpha-ketoacids (e.g., oxaloacetate), forming a feedback loop: when alpha-ketoacids accumulate, cAMP synthesis and catabolic gene expression are downregulated, allowing *E. coli* to adjust its metabolism according to nutrient availability. Given that cAMP-CRP regulates hundreds of genes^58,59^, and that catabolic proteins can represent up to 30% of the cellular proteome under certain conditions^40^ (comparable to the ribosomal sector in rich media), we reasoned that this regulator could exert effects at the global cellular scale and should not be neglected.

According to You et al.^33^, catabolic proteins form a subsector of the P sector, which is completed by the anabolic subsector. The catabolic sector strength, *β*_*C*_, can be expressed as a function of cAMP concentration, *β*_*C*_ = *β*_*C*_(*c*) (Fig. 2A). The anabolic component can be considered as passive (i.e., constitutive) to a first approximation^40^, with a constant strength *β*_*A*_. While alternative proteome partitions are possible^6,53,60^, a fundamental assumption relevant to our study is that a large part of the P sector is activated by cAMP. Therefore, we can write *β*_*P*_ = *β*_*C*_(*c*) + *β*_*A*_ = *β*_*P*_ (*c*), while for the R sector we can set *β*_*R*_ = *β*_*R*_(*g*). Thanks to Eq. (8) we obtain, at steady state,

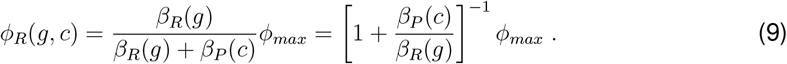

Crucially, *ϕ*_*R*_ becomes a function of both ppGpp and cAMP concentrations. The strong variation of *ϕ*_*R*_ with *g* in wild-type data can be explained in two ways: either cAMP signaling plays a minor role compared to ppGpp regulation, or cAMP levels vary jointly with ppGpp as a function of growth rate, *c* = *c*(*g*). To distinguish between these scenarios, we measured cAMP levels under the same conditions as in Fig.2B using the same methodology as for ppGpp. Consistent with the second hypothesis, we observed that ppGpp and cAMP levels covary, with their relation approximately linear (Fig.2C). As ppGpp levels increase, repressing the R sector, cAMP concentrations rise concurrently, activating the P sector. This coordinated response ensures that both cAMP and ppGpp collectively redirect resources from translation to metabolism, reflecting a joint modulation of proteome allocation by these second messengers. Importantly, our model does not assume any causal relationship between ppGpp and cAMP, nor do our data provide direct evidence for causality; for further considerations on potential interpretations, we refer to the Discussion section. We also emphasize that this positive correlation is observed specifically under catabolic limitation (C-LIM), where growth is restricted by changing the carbon source. However, our model predicts that different behaviors may emerge under other perturbations (see below).

### RelA overexpression reveals indirect regulation of ribosomal proteins by transcriptional competition with cAMP targets

Having established that ppGpp and cAMP covary positively under catabolic limitation, we reasoned that perturbations in ppGpp levels could disentangle this correlation and reveal the effects of transcriptional competition. To test this, we titrated ppGpp by overexpressing the N-terminal catalytic domain of the enzyme RelA (referred to as RelA*) using an inducible TetR-pTet system. We measured steady-state growth rate together with ppGpp and cAMP concentrations at different degrees of induction, and with different carbon sources as supplements (see STAR Methods).

Previous studies have explored RelA* overexpression and provided key insights that inform our analysis. Firstly, this perturbation is known to reduce the steady-state growth rate^37,41,61^, a finding confirmed in our data (Fig. S1A). As mentioned in the Introduction, Zhu and Dai.^37^ demonstrated that this decrease in growth rate is also associated with a reduction in rRNA content (a proxy for the ribosomal fraction^6^), while the relationship between the two (the ribosomal growth law) remains unchanged (Fig. S2A). Based on this observation, we are able to convert growth rates into ribosomal fractions with the same coefficients used for the unperturbed conditions (STAR Methods).

As mentioned in the Introduction, earlier work^41–44^ suggested that the relationship between growth rate *λ* and ppGpp *g* breaks under RelA overexpression, therefore the relationship between the ribosomal sector and ppGpp, which is central to wild-type physiology (Fig. 2B), must also be broken. This is noteworthy, as one might expect that elevating ppGpp levels in RelA over-expression mutants would simply shift cells to a slower growth regime, while still maintaining the wild-type *g* − *ϕ*_*R*_ correlation. However, our measurements confirm that the relationship between *ϕ*_*R*_ and *g* is not conserved (Fig. 3A).

**Figure 3.**
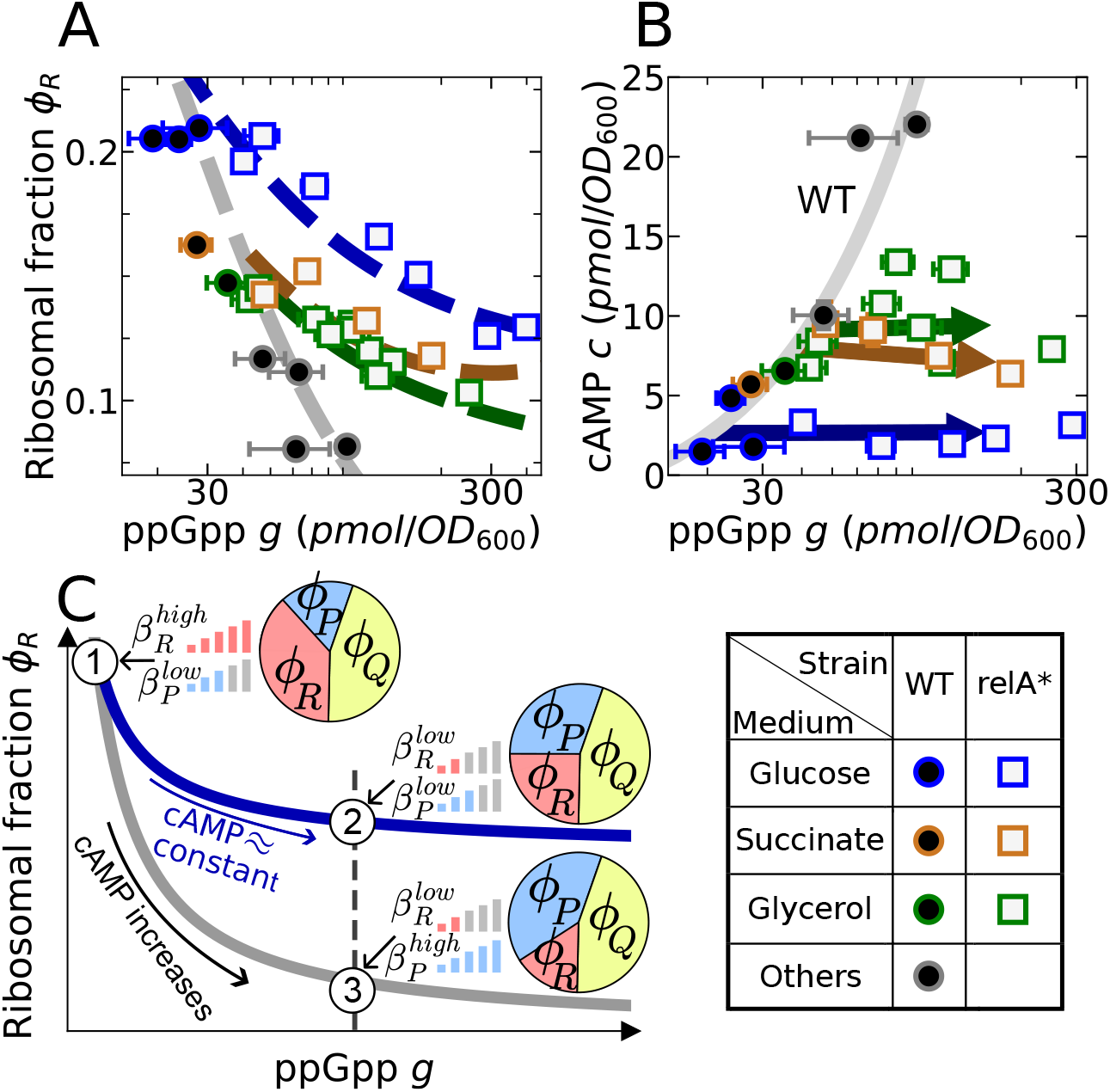
Lack of cAMP-target activation under RelA overexpression attenuates ppGpp-mediated ribosomal repression via transcriptional competition, falsifying competition-free models. (A) Upon RelA overexpression (squares), the wild-type relation between *ϕ* _*R*_ and ppGpp (circles) breaks down. By a proper choice of the functions *β* _*R*_ (*g*) and *β*_*P*_ (*c*), the model reproduces both wild-type (gray dashed line) and RelA overexpression data (colored dashed lines). To visualize the model’s predictions, we assumed that cAMP levels vary linearly with ppGpp changes during RelA overexpression (see STAR Methods), as indicated by the arrows in Fig 3B. B) The WT relation between ppGpp and cAMP (solid gray line, same as in Fig. 2C, but in log scale) is broken when over-expressing RelA. For a fixed carbon source, cAMP levels vary only minimally with increasing ppGpp. Arrows indicate the linear fits used as input for the model predictions in panel A. (C) Origin of the branching in Fig. 2A. At low ppGpp (1), ribosomal promoters operate at maximal capacity, while the P sector is minimally activated. If RelA is overexpressed starting from this condition (blue line), R is repressed by the increased ppGpp concentration, but the intrinsic activation of the P sector remains approximately constant due to unchanged cAMP levels (2). Conversely, if the same ppGpp concentration is achieved in the wild type by worsening the nutrient quality (black line), cAMP levels rise and activate the P sector. This results in a smaller ribosomal fraction in the latter case (3).

Specifically, at equivalent ppGpp concentrations, RelA* overexpression mutants, despite an overall decrease in their growth, displayed higher growth rates, and hence higher ribosomal fractions, compared to the wild type (Fig. 3A and Fig. S1A). In other words, RelA* overexpression reduces the repressive power of ppGpp on ribosome biogenesis, as the same ppGpp concentrations give higher ribosomal levels than in the unperturbed condition. Similarly, this conclusion is supported by comparing recent data from refs.^38,39^ with our wild-type data (Fig. S2C).

Thus, our observations support the idea that ribosomal content is not solely dependent on ppGpp levels. This challenges regulatory models that propose ppGpp as the exclusive regulator of proteome partitioning. Instead, our findings support a more complex scenario involving competition for RNA polymerase between multiple large-scale gene expression programs, described by our model through Eq. (9).

To further explore this deviation, we examined the relation between ppGpp and cAMP under RelA overexpression. Remarkably, cAMP concentration data points fall below, rather than on, the wild-type ppGpp-cAMP curve (Fig 3B). This implies that, at the same ppGpp concentration, the catabolic sector is less activated by cAMP compared to the physiological level of activation found in the wild-type strain. The trend in cAMP data is in line with a constant for growth in minimal medium supplemented with glucose (blue arrow), and is compatible with a constant for minimal medium supplemented with succinate (ochre arrow). For these two conditions, we can reasonably conclude that cAMP levels remain largely unaffected by RelA overexpression. The behavior observed in glycerol is more complex, yet cAMP levels still remain below the wild-type curve (gray line).

Based on these observations, we determined the functional shapes of the two sector strengths *β*_*P*_ and *β*_*R*_ required for the model to reproduce our data for both the wild type and RelA* overex-pression mutants. Using Eq. (9), we found that a model with

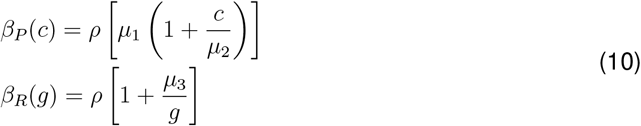

reproduces the data well (see Fig. S3A for an estimate of the goodness of fit), where *µ*_1_, *µ*_2_, *µ*_3_ can be fixed by fitting (Table S1) and *ρ* remains a free parameter. The predictions of the model can be visualized as a function of ppGpp by assuming a relationship between cAMP and ppGpp for each carbon source. Fig. 3A shows the outcome assuming the best linear fits, represented by arrows in Fig. 3B. Other representations of the cAMP-ppGpp relationship lead to slightly different visualization in the *g− ϕ*_*R*_ plane (Fig. S3B,C) without impacting the quality of the fit.

While this model is fairly rich in parameters, it gives rise to simple interpretations. The parameters *µ*_2_ and *µ*_3_ are, respectively, effective cAMP and ppGpp concentrations. The parameter *µ*_1_ accounts for different number of genes and basal promoter strengths between the two sectors. A basal rate for the R sector is necessary because extreme ppGpp overabundance does not reduce the ribosomal fraction to the lowest levels observed in the wild type; instead, *ϕ*_*R*_ appears to approach an asymptotic value, particularly for glucose and glycerol (Fig 3A, high ppGpp region). The basal rate for the P reflects the contribution from its anabolic subsector^40^ (see Discussion for details).

The model also necessarily includes the free parameter *ρ*, because our approach is only able to access the *ratio* 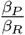 (see Eq. (9)). Therefore, the model is invariant with respect to scaling both activities by a common multiplicative factor, which potentially keeps into account transcriptional effects shared by the two sectors, as well as potential influence of ppGpp on Crp activity^62^ (Discussion and Methods S1). For simplicity, we assume the absence of such effects, treating *ρ* as a constant. However, even if these effects were present, we note that *ρ* would only affect the total promoter strength *β*_*TOT*_, without influencing competition between the R and P sectors, which is fully determined by the effective concentrations *µ*_2_ and *µ*_3_.

We can illustrate the key features of the model using glucose as a reference condition (Fig. 3C). In wild-type cells growing on glucose, ppGpp and cAMP levels are low: this leads to strong derepression of the R sector and a lack of activation of the P sector, resulting in a high ribosomal fraction, 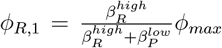. When ppGpp increases due to RelA overexpression (blue line in Fig. 3C) while keeping glucose as the carbon source, it represses the R sector. However, because cAMP levels remain constant, the P sector is not activated, leading to only a small reduction in ribosome abundance, with 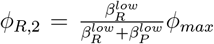. In contrast, when ppGpp increases to the same value due to worsened nutrient quality (black line in Fig. 3C), cAMP levels rise in tandem activating the P sector and resulting in a more significant downregulation of the ribosomal sector, with 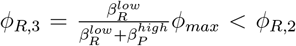. Note that this reasoning also applies when cAMP levels are not constant under RelA overexpression, as long as they remain below the wild-type cAMP-ppGpp curve, as seen in glycerol (green squares in Fig. 3B).

Thus, the relationship between ppGpp and R sector expression observed in wild-type conditions is not only a consequence of ppGpp repression but also a result of transcriptional competition (or equivalently a competition for allocation of free RNA polymerase) with catabolic genes, which are activated by cAMP. In contrast, during RelA* overexpression, the relationship between ppGpp and R sector expression reflects ppGpp repression alone, without concomitant activation of catabolism. Therefore, repression of the ribosomal sector appears weaker in this condition, as higher ppGpp levels are required to suppress ribosomal protein expression compared to wild-type conditions.

This is the central result of our study: R sector expression is determined not only by ppGpp concentrations but also by constraints imposed by RNA polymerase allocation and transcriptional competition with other sectors.

### Alternative explanatory models require transcriptional competition between sectors

To assess the robustness of our findings, we explored various alternative models to account for potential confounding factors that could contribute to the observed data. One such factor is gene dose-dependent effects, which are not explicitly included in the simplest version of our framework. While we present a method to address these in the Methods S1 document, we briefly note here that the growth rate dependence of the ori/ter ratio, which determines copy numbers, remains unchanged under ppGpp overabundance^63^. Building on this, we show that gene copy number variations do not account for the divergence in the relationship between *ϕ*_*R*_ and *g* (Methods S1). Within the examined growth rate range, these effects are minimal, and incorporating them into our fitting algorithm results only in parameter adjustments (Table 1). Moreover, we note that the branching in Fig. 3A is not caused by variations in the rate of replication initiation, which may be under complex regulation^63^. This is because our definition of regulatory functions is independent of Ori concentration, which influences total gene concentrations, but not relative levels. Additionally, it is important to emphasize that our model is independent of the speed of transcriptional elongation or cell size, variables that are both known to be affected by RelA overexpression^52,61^, but are not related to transcriptional competition.

In addition, considering the mechanisms of ppGpp-dependent repression of ribosomal genes, attention should be given to the role of DksA, which is required for ppGpp activity as a transcriptional repressor of the ribosomal sector. One might propose that ppGpp-mediated repression becomes limited by DksA under high ppGpp levels. However, data from refs.^9,64^ show that, in the wild type, DksA is maintained at a constant level across perturbations, which likely reflects negative autoregulation of its gene^65^. Therefore, we can safely exclude the possibility of a DksA-limited regime, particularly for ppGpp values in the typical wild-type range, where branching is already recorded (Fig. 3A). Indeed, existing observations report consistently high concentrations of DksA across various conditions^18^. Incidentally, the proteomics data from ref.^9^ report that the abundance of Crp, which is essential for cAMP signaling, increases under catabolic limitation in the wild type. This increase can be interpreted as an adaptation to the cellular levels of cAMP, supporting frameworks like ours^33^, where cAMP functions as a sole regulator, without explicitly considering Crp. Refined models where a Crp-cAMP regulator is limited by Crp are possible, but these details appear to go beyond the question of transcriptional competition.

In principle, other scenarios can be conceived to describe our data. For instance, one might consider allowing *ϕ*_*Q*_ to vary, leading to various model configurations. We demonstrate that a model where the sector strength *β*_*Q*_ remains unchanged during RelA overexpression, while the P sector is treated as passive, fails to explain the observed data. This alternative model predicts a greater downregulation of the R sector compared to wild-type conditions, which contradicts our findings (Methods S1 for proof). As a second alternative, we explored a model where, under RelA overexpression, proteome resources released by the R sector are allocated to the Q sector, while *ϕ*_*P*_ remains constant. Within our framework, we demonstrate that cAMP-mediated activation of the P sector is still necessary in this scenario (Methods S1). Moreover, we note that while some catabolic and anabolic genes are upregulated by ppGpp^19,66^, considering this mechanism alone without activation by cAMP would predict *ϕ*_*R*_ to depend solely on ppGpp (Methods S1). Indeed, at equivalent ppGpp concentration, the P sector would be equally activated in RelA overexpression mutants and in the wild type, imposing the same degree of (indirect) repression on *ϕ*_*R*_, contradicting the experimental findings in Fig. 3A.

Thus, we conclude that, within the range of scenarios we considered, transcriptional competition is a necessary ingredient for the model to reproduce the behavior of our experimental data.

### Suboptimal proteome allocation is maintained by distorted sensing of elongation rates

The observation that the *ϕ*_*R*_ − *λ* relationship is conserved under RelA overexpression, while the *ϕ*_*R*_ − *g* relationship is not, prompted us to investigate the behavior of other variables of the system with respect to ppGpp. Specifically, we focused on the relation between translation elongation rate *ϵ* and *g*. Wu and coworkers^20^ reported a solid link in the wild type between these two variables, even under out-of-steady-state conditions (Fig. 4A), as follows

**Figure 4.**
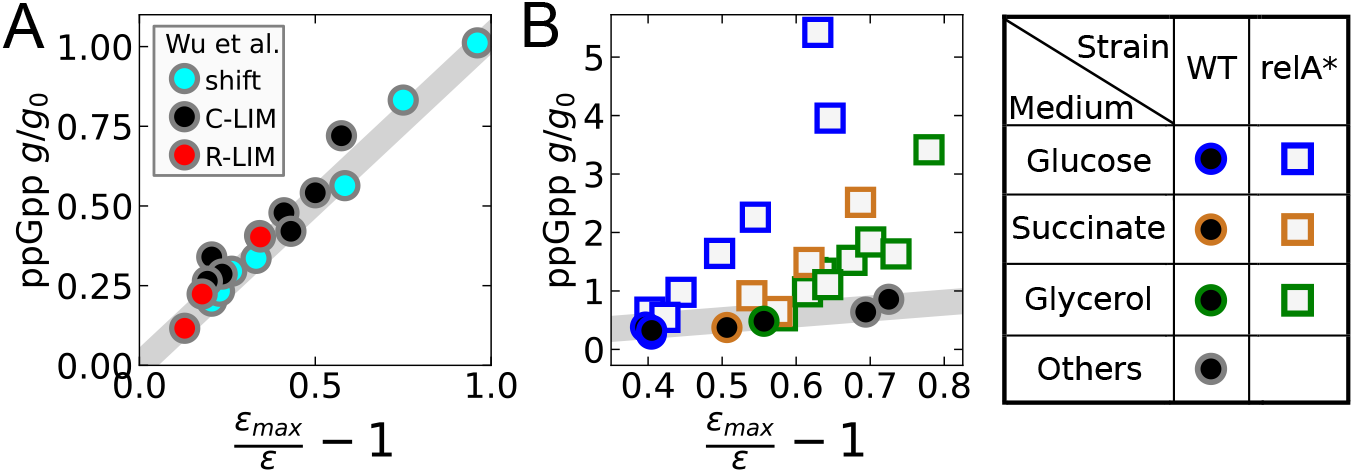
RelA* overexpression causes distorted sensing of elongation rates. A) In the wild type, ppGpp levels are determined by the elongation rate via sensing by RelA, leading to (11). Data from ref. ^20^ for a diauxic shift (light blue), catabolic limitation (black), or chloramphenicol treatment (red). B) ppGpp levels vs elongation rates as calculated from our growth rate data, using the elongation rate growth law in ref. ^37^. Our estimates show that the linear wild-type relation in panel A does not hold under RelA* overexpression. At the same elongation rate *ϵ*, ppGpp is more abundant in RelA* overexpression mutants, reflecting the excess of synthase domains and the lack of compensation by SpoT. For both panel A and panel B, gray lines represent the best linear fit of wild-type data according to (11), with *g*^0^ *≈* 70 *pmol/OD*^600^ for our data.

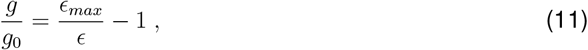

where *ϵ*_*max*_ represents the maximal elongation rate, and *g*_0_ denotes a reference ppGpp concentration. It has also been reported that the *ϵ* − *ϕ*_*R*_ (and hence *ϵ* − *λ*) relationship remains substantially unchanged under RelA* expression^37^ (Fig. S2B); therefore, ppGpp overexpression cannot preserve the relationship established by Eq. (11).

Fig. 4 explicitly shows this point using the data in ref.^37^ to convert growth rates from our experiments into elongation rate values (STAR Methods) and plot them against ppGpp concentration. As expected, Fig. 4B shows that RelA overexpression breaks Eq. (11). In fact, at equivalent elongation rates, we find higher ppGpp concentrations in RelA* overexpression mutants. This change can be described as a distorted sensing of elongation rates by the cell^20^. At equivalent values of *ϵ*, cells perceive harsher conditions than those actually present. It is important to note that if the wild type cAMP-ppGpp relation (Fig. 2C) were conserved upon RelA* overexpression, our model would predict a collapse on the wild-type curve for *ϕ*_*R*_ as a function of ppGpp, despite distorted sensing of *ϵ*. Thus, distorted elongation rate sensing, combined with disrupted cAMP-ppGpp coordination, acts to maintain suboptimal proteome allocation when RelA is overexpressed.

We also note that this prediction is compatible with the theory proposed by Wu et al.^20^ for the wild type. Here, the steady-state ppGpp concentration is given by the steady-state balance between synthesis by RelA, with rate *a*, and hydrolysis by SpoT, with first-order rate constant *b*, yielding 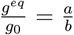. The synthesis rate is taken to be proportional to the number of “dwelling” ribosomes, and the hydrolysis rate to be proportional to the number of ribosomes in the translocating state of elongation, so that the ratio 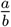 becomes a function of the elongation rate,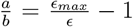, leading to (11).

Assuming this theory, we can explain ppGpp overabundance in Fig. 4 as follows. In the RelA overexpression strain, ppGpp is also synthesized by the truncated RelA enzyme, which has constitutive activity^41,67^. We write ppGpp dynamics and its equilibrium concentration as follows

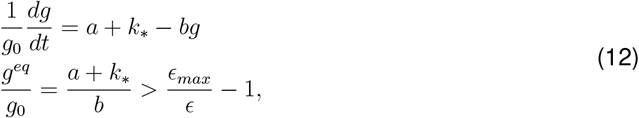

with the second equation representing the deviation in Fig. 4B. Here *k*_*\**_ represents the constitutive synthase activity of RelA*, which varies with the degree of induction. Consistent with this, we compute *k*_*\**_ from our data and demonstrate its correlation with the concentration of the pTet inducer, doxycycline (Methods S1). As a result, for the same elongation rate *ϵ* (i.e., for the same number of dwelling and translocating ribosomes), RelA* induction corresponds to increased ppGpp.

We must also note that if we focus on a single carbon source, not only is ppGpp concentration higher than in the unperturbed condition, but *ϵ* also decreases 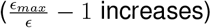. We attribute this change to the feedback loop between ppGpp and the translation apparatus^36,46^. We expect increasing ppGpp through RelA overexpression and keeping the nutrient source fixed leads to transcriptional repression of tRNAs and translation factors, which are co-regulated with ribosomes. According to the theory by Dai et al.^47^, this would in turn cause *ϵ* to decrease.

### Transcriptional regulation of hibernation factors explains deviations in ribosome inactivity

In addition to elongation rate *ϵ* and ribosome proteome fraction *ϕ*_*R*_, the fraction of actively translating ribosomes, *f*_*a*_, is a crucial determinant of growth rate^47^. We can calculate *f*_*a*_ using the relationship 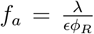. This definition assumes negligible protein degradation, which plays a role in setting the fraction of active ribosomes, particularly at very low growth rates^48^. In the following, we maintain this simplifying assumption, as it provides a reasonable approximation in moderate-to-fast growth conditions^36,48^.

The main contribution to inactive ribosomes is generally attributed to ribosome deactivation by hibernation factors^20^. Other factors, such as stalled ribosomes^47^ and mRNA scarcity^29^ might also play a role, but we focus for simplicity on inactivation by hibernation factors, following the model proposed in ref.^20^. In *E. coli*, four main hibernation factors, Rmf, RaiA, RsfS, and Hpf, dimerize or sequester ribosomes and their subunits^68–70^, thereby preventing them from initiating translation (Fig. 5A). We group these factors into a new sector characterized by a mass fraction *ϕ*_*HIB*_ and assume complete binding of hibernation factors to ribosomes, expressed as

**Figure 5.**
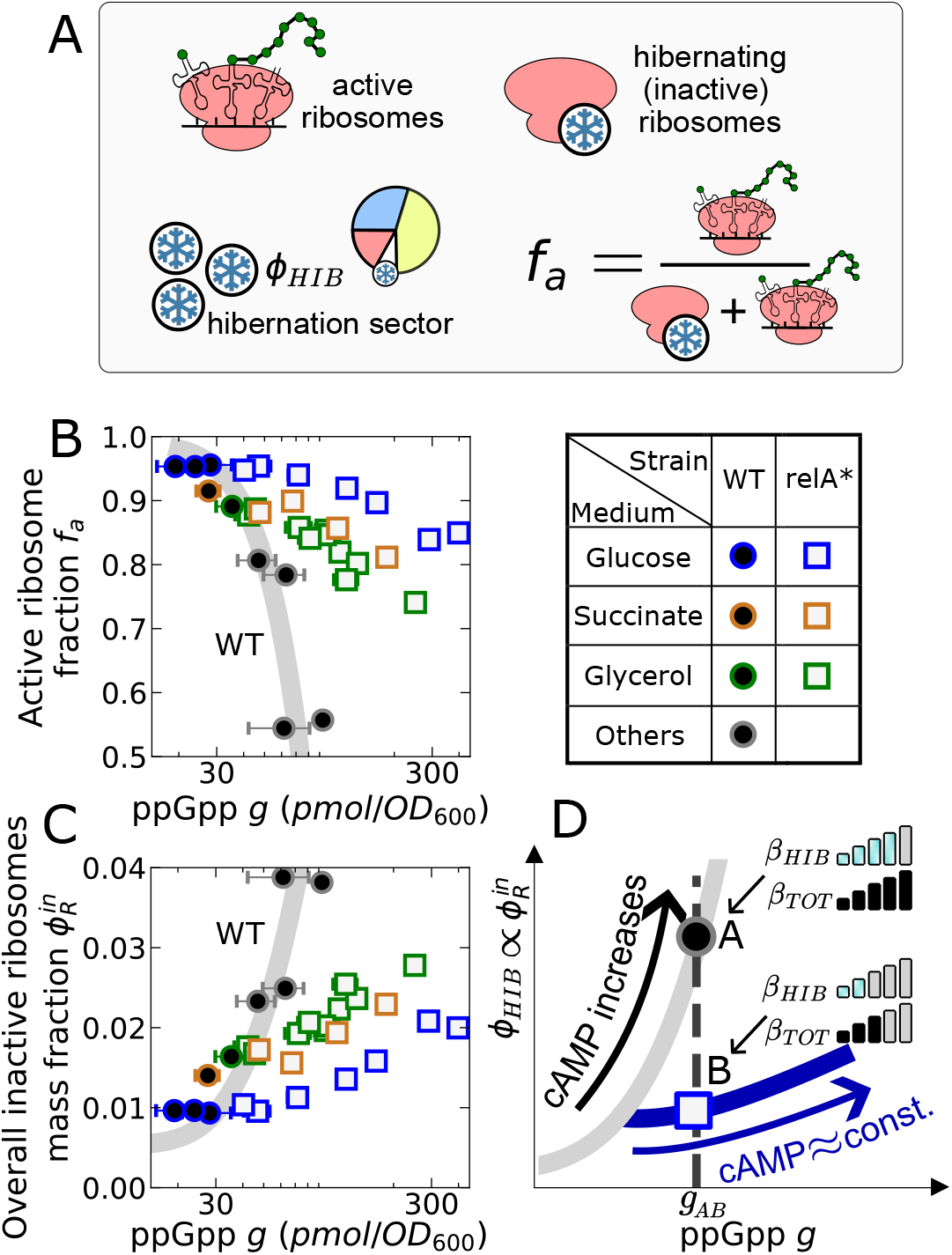
Transcriptional activation of hibernation factors by cAMP explains deviations in the fraction of active ribosomes. (A) Scheme of the model. We consider two populations of ribosomes: actively elongating ribosomes (with fraction *f*^*a*^), which contribute to translation and growth, and hibernating (inactive) ribosomes. The proteome sector containing the hibernation factors has mass fraction *ϕ* ^*HIB*^. (B) Estimates for the fraction of active ribosomes 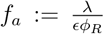, calculated from our growth rate data, plotted against ppGpp. The gray line is obtained with this definition, using the wild-type growth laws for *ϵ* and *ϕ* ^*R*^ as in ref. ^47^ and ref. ^37^ (see STAR Methods). Under RelA overexpression, *f*^*a*^ decreases as a function of ppGpp, but its decrease is less pronounced with respect to unperturbed data. (C) Under RelA overexpression, the estimated proteome fraction corresponding to inactive ribosomes 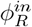 increases more gradually with ppGpp. The gray wild-type line is obtained analogously to the one in panel A, through equation (13) (STAR Methods).(D) Results in panel B can be explained by a transcriptional competition theory where hibernation factors are activated by both ppGpp and cAMP, with a strength *β* ^*HIB*^, explaining their scarcity under RelA overexpression.

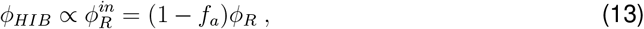

again in line with Wu et al.^20^. Here 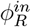 is the mass fraction associated with the inactive ribosomal sector.

Knowledge of *λ* and the values of *ϕ*_*R*_ and *ϵ* allow us to calculate *f*_*a*_ and 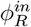 in our data and plot them against ppGpp concentration (Fig 5B and 5C). Our results reveal a significant deviation from the wild-type relationship: at equivalent ppGpp concentrations, RelA overexpression mutants display higher values of the active fraction *f*_*a*_, despite an overall decrease with over-expression. This suggests that *ϕ*_*HIB*_ is not regulated exclusively by ppGpp, as proposed in the model of ref.^20^.

We speculated that this fact might also be a consequence of transcriptional competition. To explore this possibility, we introduce a collective promoter strength associated with hibernation factors, *β*_*HIB*_. Fig. 5D provides a schematic representation of how second messengers could be influencing this variable in relation with the mass fraction *ϕ*_*HIB*_, taking again glucose as a reference. At the same ppGpp concentration *g*_*AB*_, the wild type (point A) presents a higher cAMP concentration than the RelA overexpression mutant (point B), i.e., *c*_*A*_ *> c*_*B*_. Our data indicate that *ϕ*_*HIB*_(*B*) *> ϕ*_*HIB*_(*A*), which at steady-state can be written in terms of promoter strengths from (7) (STAR methods for derivation)

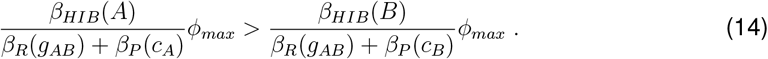

This inequality captures the competition between the hibernation sector in the numerators and the P and R sectors in the denominators. Since the denominator is greater for the wild type (left-hand side) due to the activation of the P sector by cAMP, the inequality suggests that the strength of the hibernation sector must also be higher in wild-type cells, i.e., *β*_*HIB*_(*A*) *> β*_*HIB*_(*B*).

This model shows that the data are consistent with cAMP activating the hibernation sector, in addition with the reported effect by ppGpp^20^. Notably, here the deviation from the wild-type line arises from a direct regulatory effect, rather than simply from competition for RNA polymerase, as was observed for the ribosomal fraction *ϕ*_*R*_ (Fig. 3C). Indeed, global coupling acts on the denominator, and taken alone it would cause the opposite relationship: *ϕ*_*HIB*_(*A*) *< ϕ*_*HIB*_(*B*) due to activation of the P sector by cAMP, with *β*_*P*_ (*c*_*A*_) *> β*_*P*_ (*c*_*B*_). Therefore, we hypothesize that cAMP may regulate a direct transcriptional activation of the hibernation sector. In line with this claim, cAMP was found to activate transcription of *rmf* and *raiA*^70,71^.

This transcriptional model for hibernation factors may also explain why the ribosomal growth law *ϕ*_*R*_(*λ*) remains the same of the wild type under anabolic limitation^33,40^ and in the “cAMP titration” strain examined in ref.^40^. Its conservation suggests that both *ϵ* and *f*_*a*_ must are able to adjust to ribosomal abundance across these perturbations, which we predict to break the ppGpp-cAMP relationship seen under catabolic limitation (see below).

### The model predicts ppGpp–ribosome divergence under alternative perturbations

In addition to RelA overexpression, there may be other situations where competition between regulons determines a deviation from the ppGpp-ribosome relationship observed under C-LIM. An important example is titration of the enzyme glutamate synthase^40^, which imposes an anabolic limitation (A-LIM). In this condition, cAMP levels decrease as growth rate declines, and ribosome abundance falls according to the same growth law observed under C-LIM^33,40^. The catabolic sector also decreases, which can be attributed to regulation by cAMP. However, indirect effects of cAMP alone would be expected to increase the ribosomal fraction through transcriptional competition. Therefore, the observed reduction in ribosomes must reflect direct repression by elevated ppGpp^1^. Thus, in the light of our model, these findings support the interpretation that ppGpp and cAMP should be anticorrelated under A-LIM^40^. In other words, if this perturbation originates from a point on the C-LIM curve, the system should move into the region of the ppGpp–cAMP plane below that curve (Fig. 6, solid green arrow). Based on this interpretation, we predict indirect cAMP effects will create an apparent weakening of ppGpp’s repressive power (Methods S1), similar to what we observed under RelA overexpression (Fig. 3). Thus, transcriptional competition effects should be detectable as a divergence from the C-LIM relationship between ppGpp and *ϕ*_*R*_, as indicated by the dashed green arrow in Fig. 6.

**Figure 6.**
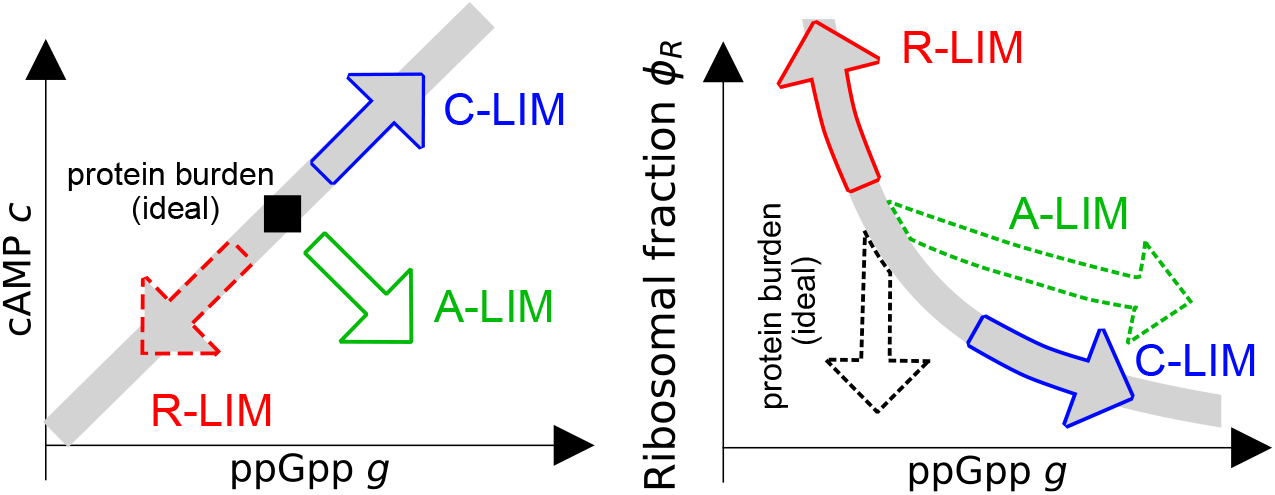
The C-LIM ppGpp-ribosome relationship can be preserved or broken under different perturbations. In our framework, different perturbations can be visualized as displacements in the ppGpp-cAMP plane. Following the C-LIM curve in this plane (gray line, left) is both necessary and sufficient for a perturbation to preserve the C-LIM ppGpp-ribosome relationship (gray line, right), due to equation (9). This principle allows us to make predictions (dashed arrows) for untested perturbations based on relationships inferred from existing literature measurements (solid arrows). Under anabolic limitation (A-LIM, green), assuming that ppGpp and cAMP anticorrelate^40^ leads to a divergence in the ppGpp-ribosome plane. We predict ppGpp and cAMP should covary under chloramphenicol treatment (R-LIM, red), as literature data^20,47^ suggest the ppGpp-ribosome relationship should be conserved. Expression of a useless protein (black) can ideally lead to a decrease in ribosome content without any change in ppGpp, due to RNAP diversion.

Another situation where competition could emerge is growth inhibition by antibiotics such as chloramphenicol (R-LIM), which leads to ppGpp depletion, opposite to RelA overexpression^6,9,47^. These two perturbations are fundamentally different in that chloramphenicol leaves the feedback loop between ppGpp and translation machinery intact. Indeed, it appears to preserve the key regulatory relationship between ppGpp and elongation rate^20^ (Fig. 4A), as well as the one between elongation rate and ribosomal sector *ϕ*_*R*_ ^47^ (Fig. S4). Given these conserved links, we conclude that the ppGpp–*ϕ*_*R*_ relationship should also be conserved, as shown by the solid red arrow in Fig. 6. (Methods S1). While paired ppGpp–cAMP measurements are not available under this condition, the ppGpp-*ϕ*_*R*_ conservation emerges in the model if cAMP and ppGpp covary (dashed red arrow in Fig. 6). Consistent with this inference, chloramphenicol has been reported to interfere with cAMP signaling^72,73^; however, it also induces distortions that require careful consideration (see Discussion).

Lastly, we consider a perturbation by overexpression of a useless protein^6,33^. Here, the details of ppGpp regulation (e.g., whether the *ϵ− g* relationship in Fig. 4A is maintained) remain incompletely understood, and no joint ppGpp–cAMP measurements are available. Therefore, we examine an idealized scenario in which overexpression does not alter ppGpp or cAMP levels. Even in the absence of such changes, allocation of RNAP to express the useless gene can decrease the ribosomal fraction (dashed black arrow in Fig. 6) via transcriptional competition (Methods S1). Thus, we predict that, in this simplified scenario, transcriptional competition should manifest again as a splitting in the ribosome–ppGpp relationship from the C-LIM line.

## DISCUSSION

While there is a large body of work on how different regulators control the various components of the proteome^20,33,40,57^, the impact of these regulatory circuits on the global scale is relatively unexplored^1,31,40,74–76^. In *Escherichia coli*, the prevailing consensus sees ppGpp alone as a governor of ribosome allocation, overriding the influence of other regulators^16,40^. However, data collected in this study confirm the previous observation^41,42,44^ that ribosome allocation is not solely dependent on ppGpp outside physiological conditions. Indeed, ppGpp’s capacity to repress expression of ribosomes is decreased under RelA overexpression (Fig. 3A).

The primary result of our combined approach integrating coarse-grained modeling and data is that allocation of RNA polymerase towards catabolic genes decreases expression of ribosomal genes. This is a form of indirect repression that requires no direct regulatory interactions between Crp and cAMP with ribosomal promoters. Instead, cAMP acts by inducing the expression of catabolic genes, which compete for RNAP allocation with ribosomal protein genes (Fig. 3C). RelA overexpression was required to reveal this competition, which is otherwise obscured in the wild type by the approximately linear relationship between cAMP and ppGpp (Fig. 2C).

This relationship, observed under catabolic limitation, is consistent with theoretical predictions by Erickson et al.^53^ and Scott and Hwa^1^. They propose that the correlation between ppGpp and cAMP arises from a mechanistic link between the two messengers, possibly due to their sensitivity to ketoacids and amino acids, which are coupled via transamination. Under this framework, the disruption of the correlation under RelA overexpression could be attributed to ppGpp’s distorted sensing of the elongation rate and, consequently, of the amino acid pool. Alternatively, the correlation might be non-causal, simply reflecting the shared role of ppGpp and cAMP as stress signals under catabolic limitation. Whether this relationship is causal remains an open question for future investigation.

Our theory places cAMP on par with ppGpp as “orthogonal” signals: not only do they control different parts of the proteome, but their levels must also be independently adjusted, since we observed deviations from the wild-type ppGpp-cAMP curve under RelA overexpression (Fig. 3B). However, some interactions have been reported, ranging from ppGpp-induced acetylation of Crp^77^ and direct binding of ppGpp to Crp, which can impair its activity^62,78^. These non-orthogonal effects can be included in the *ρ* factor in (9), without substantial changes to the model (Methods S1).

The anabolic (A) sector plays a key role in the interplay between ppGpp and cAMP, as it links catabolism to ribosomes^1,33^. In our work, we approximated it as constitutive (passively regulated), following the approach of Kochanowski and coworkers.^40^. In line with their conclusion, we emphasize that this is only an approximation, and pathway-specific regulators may dominate the control of this sector. Indeed, downregulation of the A sector under catabolic limitation^9,33^, as observed experimentally, is inconsistent with constitutive-like behavior, since expression from constitutive promoters is expected to increase in this condition^31^. To demonstrate that our model remains valid regardless of this approximation, we have attempted a more data-driven approach in defining the sector that competes with ribosomes, using gene targets of cAMP as cataloged in databases such as RegulonDB^58^ and EcoCyc^59^ (consulted February 2025), rather than relying on phenomenological definitions of proteome sectors. Our analysis revealed that cAMP-activated genes are distributed across the entire proteome, including the Q sector (Fig. S5A,B). As a result, ribosomal proteins may be considered in competition for RNA polymerase with a “cAMP-activated sector” that does not perfectly overlap with catabolic proteins. This variant adequately describes the data, requiring only a modification of parameters (Table 1 and Supplemental Note).

One important additional layer of regulation, particularly under conditions of high ppGpp, is the contribution of alternative sigma factors^79^, which can be naturally incorporated into our model (see Methods S1 for mathematical details). Alternative sigma factor-induced genes can compete with RpoD-dependent genes in several ways. First, as classically noted, they may compete for core RNAP (the well-known sigma factor competition^79,80^), a scenario that is unlikely to be quantitatively relevant in moderate-to-fast-growing wild type, given that core RNAP is typically in excess and maintained at a constant level^31^. Second, mRNAs expressed from alternative sigmulons can reduce the fraction of the transcriptome corresponding to RpoD-dependent genes, thereby decreasing their protein mass fraction, without affecting their total mRNA abundance. This effect can be captured by the model in the main text (equation (7)), simply by introducing a sector corresponding to the alternative sigmulon (Methods S1).

Although alternative sigma factors may respond to second messengers^79,81^, their quantita-tive cross-talk remains unclear. In principle, they could become relevant under extreme ppGpp overabundance, where cellular adaptations to growth slowdowns, which fall outside the scope of our present study, could influence the system. However, recent data by Zhu et al.^39^ show that, although RelA overexpression in glucose upregulates RpoS-dependent genes, the overall response remains largely unaltered in an *rpoS* knockout. Furthermore, our proteomic analysis of the sigma factor landscape under RelA overexpression (based on data from Zhu et al.^38,39^) indicates that the other sigma factors remain substantially unresponsive, and core RNAP is maintained in excess (Fig. S6), despite a fivefold increase in ppGpp. Analysis of the same dataset revealed that RelA overexpression activates cAMP-responsive genes (Fig. S5B), consistent with predictions from our model via indirect effects (Figure 3C). Overall, these observations indicate that alternative sigma factors do not strongly affect the system under the conditions examined; nonetheless, their interplay with global transcriptional programs remains to be explored.

Our model can be applied to predict responses for perturbations beyond RelA overexpression (Fig. 6). We predict that, under anabolic limitation (A-LIM), the negative correlation between cAMP and ppGpp should cause a divergence from the C-LIM ppGpp-*ϕ*_*R*_ relationship. In future studies, joint measurements of ppGpp and ribosome abundance under A-LIM could provide further model validation and allow quantification of transcriptional competition in this condition. Conversely, based on literature data and our model, we infer that chloramphenicol treatment (R-LIM) should preserve the cAMP-ppGpp covariance. We note, however, that chloramphenicol can induce polarity effects due to premature transcriptional termination in ribosomal protein operons, causing a progressive decrease in mRNA abundance along the operon^37^. As a result, the assumption of a single “coarse-grained” initiation rate for all ribosomal genes may be compromised, as ribosomal mRNA production becomes non-stoichiometric. Therefore, testing the model under chloramphenicol would require not only measuring growth rate or ribosome abundance but also accounting for the distortive effects introduced by polarity.

The remarkably robust conservation of the ribosomal growth law *ϕ*_*R*_(*λ*) under both ppGpp and cAMP titration^37,40^ under lack of coupling between the two messengers requires an explanation. To maintain consistency, we showed that the fraction of active ribosomes *f*_*a*_ must not depend solely on ppGpp. Our analysis shows that a theory of transcriptional regulation of ribosome-hibernating factors can capture this behaviour. Specifically, we speculate that activation of the hibernation sector by cAMP could be one of the factors contributing to this observation, thereby recreating the growth law. While there is evidence of a role of cAMP in regulating this sector^70,71^, additional studies on this point are required, as hibernation factors could also be regulated post-transcriptionally (raiA and rmf in ref.^82^), or additional mechanisms might influence their binding to ribosomes^70^.

More widely, our findings show how the global effects of gene transcription should be understood within a framework of RNAP allocation^30,31,83^, similar to the established proteome allocation model. Rather than viewing ppGpp as a direct regulator of ribosomal content, we show that its effects arise through global RNAP redistribution, influenced by competition with other transcriptional programs. This perspective clarifies why changes in ppGpp levels alone do not fully predict ribosome synthesis and why conditions like ppGpp overabundance alter proteome composition in ways not captured by traditional promoter-centric models. Although second-messenger signaling varies across organisms (for example, catabolite repression and ppGpp function differently in other bacteria^84–86^), our framework provides a generalizable approach for understanding global competition effects in transcriptional regulation. As a result, we believe that the principles and consequences of competition described in this study may play a general role in cellular responses across different species.

## RESOURCE AVAILABILITY

### Lead contact

Requests for further information and resources should be directed to and will be fulfilled by the lead contact, Marco Cosentino Lagomarsino.

### Materials availability

This study did not generate new materials.

### Data and code availability

- Original code and data are available on GitHub (see Key Resources Table)

## ACKNOWLEDGMENTS

We thank Gabriele Micali, Luca Ciandrini, Bianca Sclavi, Philippe Fuchs, Martina Dal Bello and Paul Wiggins for useful discussions on this work. We thank Ferhat Büke, Marek Noga, Flora Yang, and Adja Zoumaro-Djayoon for technical assistance with ppGpp and cAMP quantification. The research leading to these results has received funding from AIRC under IG 2024 - ID. 30391 project – P.I. Marco Cosentino Lagomarsino. R.D. was supported by the AIRC fellowship Italy Pre-Doc 2022 (ID 28176). G.B. is supported by the Netherlands Organization for Scientific Research (NWO). M.L. was supported as part of the Frontiers of Nanoscience program.

## AUTHOR CONTRIBUTIONS

Conceptualization, G.B. and M.C.L.; theoretical investigation, A.R. and R.D.; experiments, M.L.; data analysis, A.R., M.L., and R.D; writing A.R. and M.C.L.; funding acquisition, M.C.L., G.B., and R.D.; supervision, M.C.L. and G.B.

## DECLARATION OF GENERATIVE AI AND AI-ASSISTED TECHNOLOGIES

During the preparation of this work, the authors used ChatGPT for minor rephrasing of the text. After using this tool, the authors reviewed and edited the content as needed and take full responsibility for the content of the publication.

## SUPPLEMENTAL INFORMATION INDEX

### Document S1

Figures S1-S6 and their legends, Table S1.

### Methods S1

Detailed information about modeling.

## STAR METHODS

### Key Resources Table

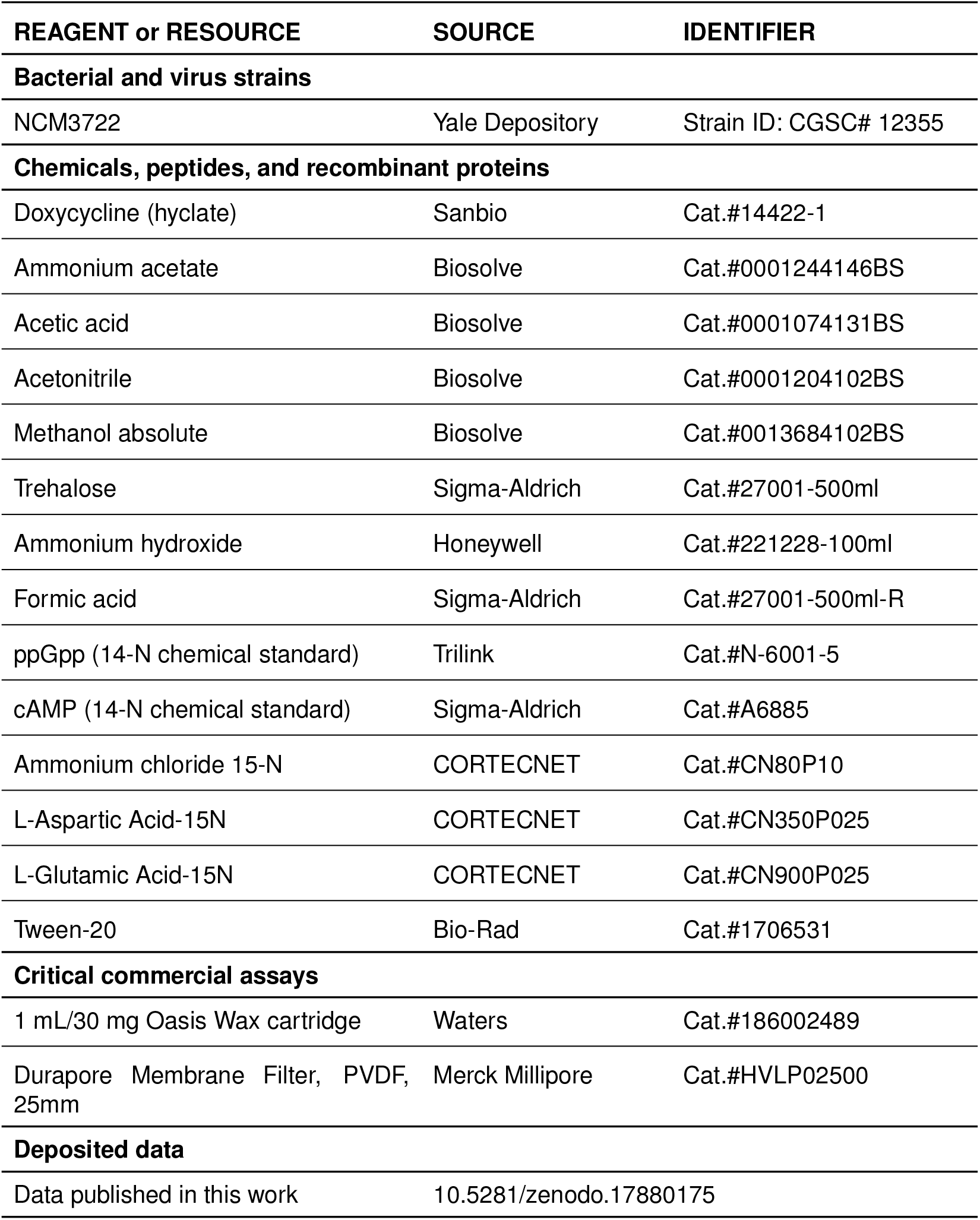
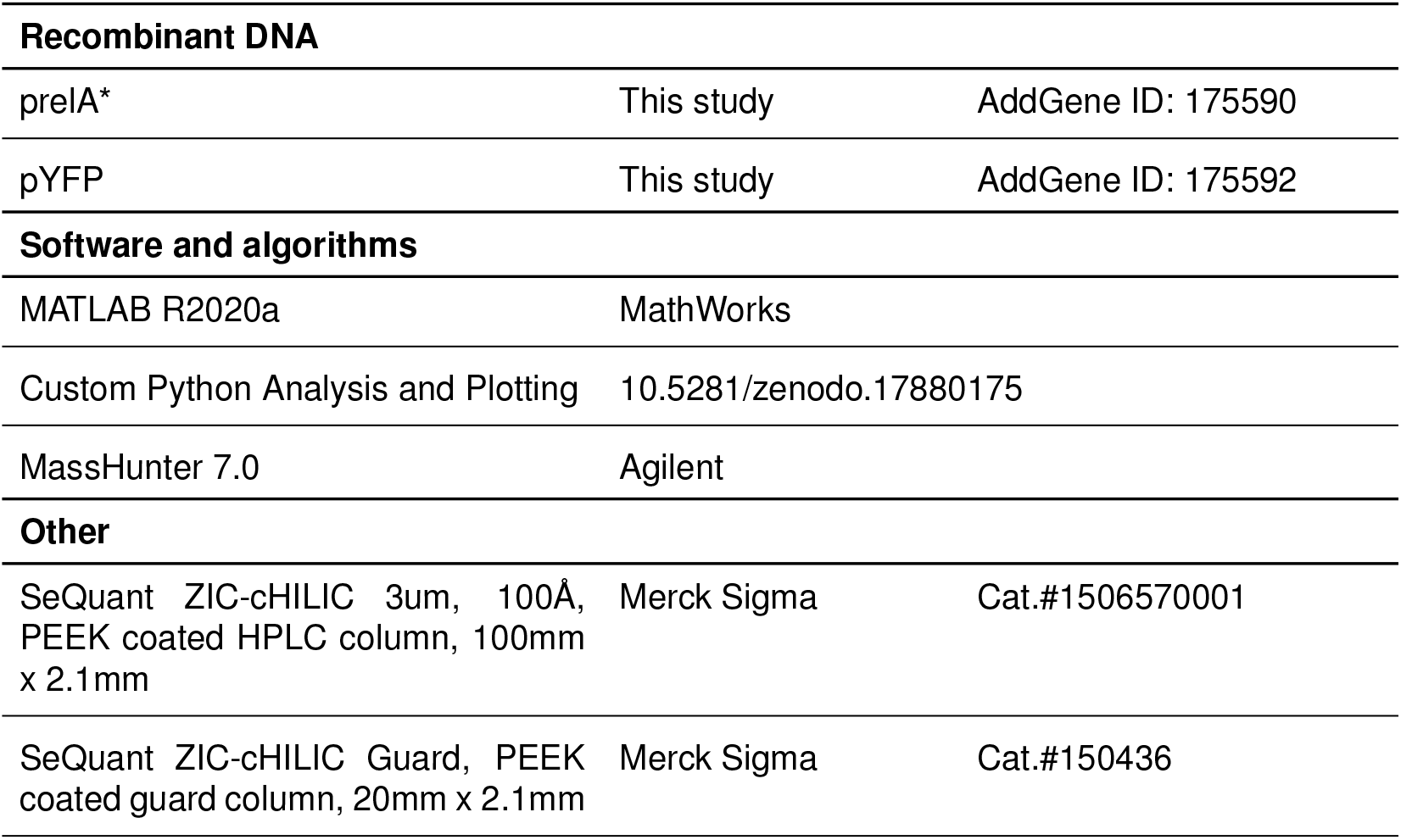

### Experimental model details

#### Strains and plasmids

All experiments were performed using Escherichia coli wild-type strain NCM3722 (CGSC# 12355) and its derivative carrying a RelA* overexpression plasmid (pRelA*). pRelA* (Tet promoter, kanamycin resistance, sc101** origin) encodes a fusion of the N-terminal 455 amino acids of the E. coli RelA with mVenus as described in Buke et al^61^ and is available from the Addgene plasmid repository (Plasmid #175590).

#### Culture conditions

Cultures were maintained as described in Noga et al.^42^ Cells were inoculated from a LB agar plate into 3 mL overnight cultures of the corresponding medium to be used the next day. Main cultures were prepared in 250 mL Erlenmeyer flasks containing at least 25 mL of fresh MOPS minimal medium supplemented with 0.2% (w/v) carbon source, which could be glucose, succinate, glycerol, acetate, aspartate or glutamate. Cultures with the pRelA* plasmid also contained 25 µg/mL kanamycin. (See resource table for reagents list.) Main cultures were inoculated from overnight precultures to 1/100 of the culture volume. Flasks were maintained at 37 °C in a Grant Instruments Sub Aqua Pro dual water bath and contained a 12-mm-long magnetic stir bar (VWR) coupled to a submerged magnetic stir plate (2mag MIXdrive 1 Eco and MIXcontrol 20) set at 300 rpm. To enable discrimination of cellular ppGpp and cAMP from the internal standard, the medium contained isotopically labeled ^15^NH_4_Cl or ^15^N labeled aspartate or glutamate as nitrogen sources.

### Method details

#### Sample collection and analysis for ppGpp and cAMP quantification

Samples were collected and processed largely as described in Buke et al.^61^ For ppGpp titration, bulk cultures were induced with doxycycline at an optical density (600 nm) of between 0.05-0.08 (early exponential phase). For each condition, three sampling replicates were collected. Samples were harvested at OD *∼*0.35 for wild-type strains, while samples from induced pRelA* cultures were sampled only after growth rate stabilized (steady-state was defined as a variation of less than 10% in growth rate over a 30-minute period), which required between 3-5 hours. In all cases samples were taken between 0.3 and 0.4 OD. Three technical replicates were sampled by pipetting 1mL or 2.5mL (for lower ppGpp concentrations) of culture onto a wet filter under vacuum. The filter was immediately quenched upside down in 1mL ice-cold 2M formic acid. After filtering, a known concentration of the nucleotides of interest was added to the quenching solution. After 30 minutes in this solution, the filter was thoroughly washed and discarded. 25*µ*L ammonium hydroxyde was added to the collected quenching solution in a 1mL tube which was frozen at -80 °C and was kept at least overnight and up to one month.

#### Growth rate measurements

Growth rates were quantified by continuously monitoring culture turbidity with an automated absorbance measurement system using a spectrophotometer (SPECTROstar Nano, BMG Labtech). Liquid culture from the 37 °C water bath flasks was continuously circulated through a flow-through cuvette (Starna Cells spectrophotometer flow cuvette 583.4-Q-10/Z8.5) using a peristaltic pump (Fisher Scientific GP1000). Optical density was measured every 5 s to 1 min, depending upon the growth rate. Growth rates were calculated by smoothing OD measurements with a 5-min wide Gaussian filter and computing the median of the time derivative of the natural logarithm of OD over a 1-h sliding window to precisely determine steady-state growth rates.

#### ppGpp and cAMP quantification via LC/MS

To measure nucleotides, samples were thawed on ice, then sonicated in ice-cold water for 10 minutes. Solid phase extraction was performed to purify nucleotides and get rid of other compounds that might cause matrix effects on the compounds of interest by altering the chromatographic separation or suppressing their ionization. To do so, during sonication, an Oasis Wax SPE cartridge 30mg 30µm was prepared by, first, flowing through three times 1mL methanol, then, three times 1mL 4.5 pH 50mM ammonium acetate. The sample was then applied on the cartridge and slowly flowed through. The cartridge was washed with 1mL 4.5 pH 50mM ammonium acetate followed by 1mL methanol and left to dry under vacuum for 5 minutes. The sample was eluted very slowly from the cartridge into an Eppendorf tube with 200µL 5:3:1:1 MeOH:ACN:H2O:NH4OH. The obtained sample was concentrated using a vacuum centrifuge after addition of 10µL of 5% trehalose for stabilization during drying. Once dried, the sample pellet was rediluted in 20µL 5:3:2 MeOH:ACN:H2O, then analyzed with an Agilent triple quad mass spectrometer and a ZIC-cHILIC column. For the first minute we maintained 90% mobile phase B 11.25mM ammonium acetate, 3.75mM acetic acid and 2mM acetylacetone in 80% ACN and 10% mobile phase A 3.75mM ammonium acetate, 1.25mM acetic acid and 2mM acetylacetone in water. Then we applied a gradient towards reaching 80% mobile phase A at 15 minutes. We maintained this concentration for one minute before applying a new gradient towards 100% mobile phase B between 16 and 18 minutes. We maintained this concentration until 22.5 minutes to flush out remaining compounds. For each compound, 14N and 15N peaks were obtained, corresponding respectively to the added standard concentration and the biological sample. Concentrations in pmol/OD600 were obtained by multiplying the 15N/14N area ratio by the added compound concentration, then normalizing by OD600 at sampling. The OD600 value was determined as the average of the last measurement before sampling and the first measurement after sampling. Error bars represent standard errors reflecting variability across technical replicates.

### Quantification and statistical analysis

#### Conversion of growth rates into ribosomal fractions and elongation rates

To convert our growth rate data (Fig S1) into ribosomal fractions, we applied a linear fit to the total RNA/protein ratio measurements *r* from Zhu et al.^37^, leveraging the conservation of the ribosomal growth law (Fig S2A):

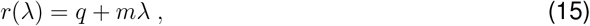

with *q* = 0.073 and *m* = 0.217 *h*. We note that interpolating data from Dai et al.^47^ yields substantially the same coefficients *q* and *m*. To obtain the “extended” ribosomal fraction, we then transformed RNA/protein ratios with the following conversion:

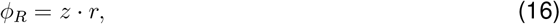

with *𝓏* = 0.76, as discussed in Scott et al.^6^.

To convert our growth rate data into elongation rate values, we leveraged the conservation of the elongation rate growth law under RelA overexpression^37^ (Fig.S2B) and computed

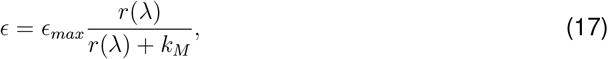

with *ϵ*_*max*_ = 6.44 *h*^*−*1^ (equivalent to 22 amino acids/s, see Methods S1) and *k*_*M*_ = 0.11 as in Dai et al.^47^. Note that Fig. 4 is independent of the value we choose for *ϵ*_*max*_, as only the ratio 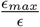 appears in the definition of the x-axis.

The knowledge of the relationships *ϵ*(*λ*) and *ϕ*_*R*_(*λ*) allows us to compute 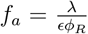 for each data point in this study.

#### Generating the wild-type active ribosomes-ppGpp curve (Fig. 5)

To generate the wild-type gray curves in Fig. 5B and Fig. 5C, we first fitted a linear relationship between *ϕ*_*R*_ and ppGpp (following ref.^20^), 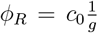, as in Fig. 2B. The quantities *λ* and *ϵ* can be obtained from *ϕ*_*R*_ by inverting the relationships (15), (16) and (17) found in the previous paragraph. This allows us to express 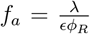 as a function of ppGpp (Fig. 5B). The wild-type curve in Fig. 5C can be computed from *f*_*a*_ and *ϕ*_*R*_ through the relation 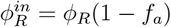.

#### Modeling - coarse grained promoter strengths

To define the coarse-grained sector strengths in equation (5), we start from (3), and sum over the genes belonging to the same sector, leading to the coarse-grained messenger dynamics

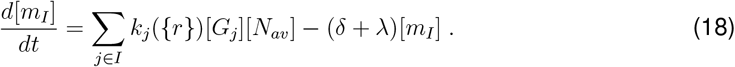

We define the coarse-grained “sector strengths”, as

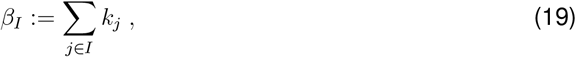

which gives (5) if all genes are assumed to be present at the same concentration [*G*].

#### Modeling - gene dosage effects

To incorporate gene dosage effects in our model, we express gene concentrations relative to the origin of replication (Ori), [*G*_*i*_] = *d*_*i*_(*λ, x*_*i*_)[*Ori*], with the dose *d*_*i*_ depending on the normalized distance *x*_*i*_ between gene and Ori, with the replication terminus (Ter) corresponding to *x* = 1^31,87,88^. For each sector, we can define a “regulatory activity”, consistently with the terminology of Balakrishnan et al..^31^, as

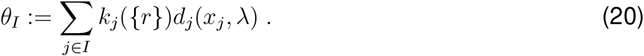

Using this definition, we obtain an equation describing the transcripts dynamics for each sector,

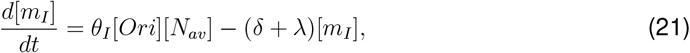

which incorporates both transcription factor regulation and gene dosage effects.

In fast growth conditions, the relative abundance of genes located near the origin of replication tends to increase, enhancing the expression of certain genes, such as those encoding ribosomal proteins and ribosomal RNA^89^. To include this in our model, we use a mean-field approximation by assigning each gene in a given sector to a single chromosomal position *x*_*I*_, thus writing 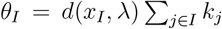. To obtain the parameters in Table 1, we set *x*_*R*_ = 0.25, and *x*_*P*_ = 0.6, reflecting the advantage at fast growth for ribosomal genes (see Methods S1 for details on how these values were estimated). This approximation leads to the factorization

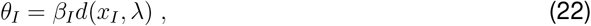

which separates the influence of transcription factors from gene dosage effects. With this setting, one still finds (6), but with the definition

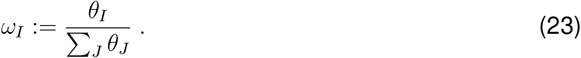

Methods S1 provides the full mathematical formulation of the model variant that includes gene dosage effects, which we used to replicate the data analyses and parameter fitting described in the main text. As already described by Balakrishnan et al.^31^, gene dosage effects are small in the range of conditions explored in this study (0.2 *−*1 *h*^*−*1^). This model variant can be mapped onto the simplified version used in the main text by assuming the same gene copy number for all sectors (e.g., by setting *x*_*I*_ = 0.5 for all *I*). This leads to:

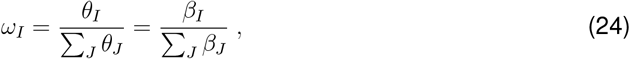

which then gives (7) in the steady state.

#### Fitting the model of transcriptional competition to ribosomal fractions

Data fitting of ribosomal fractions was performed through Orthogonal Distance Regression. A detailed explanation of the fitting approach can be found in Methods S1. Although we did not perform comparative modeling, we can assess the goodness of fit for our model in Fig. 3A by calculating a coefficient of determination. Specifically, we compared a linear model that uses our growth rate prediction (based on ppGpp and cAMP concentrations for each condition as inputs) as the explanatory variable against the measured growth rate. The results are shown in Table 1, and visualized in Fig. S3A for the model used in the main text (without gene dosage effects).

#### Transcriptional competition theory for hibernation factors

To explain the deviations in *f*_*a*_ as a function of ppGpp (Fig. 5), we focused on ribosome hibernation factors. We group these factors into a new sector characterized by a mass fraction *ϕ*_*HIB*_ and a regulatory activity *θ*_*HIB*_. We assume that hibernation factors are in competition with P and R, and use the same *ϕ*_*Q*_ and *ϕ*_*max*_ of the three sector model: equation 7 for the hibernation sector reads

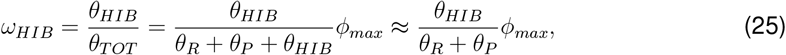

where we neglect the term *θ*_*HIB*_ at the denominator given the small size of the hibernation sector, based on the proteomics data in ref.^9^. At steady-state *ω*_*HIB*_ = *ϕ*_*HIB*_ due to (7), and ignoring gene dosage effects, we find from the above equation

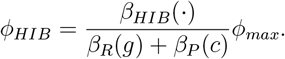

Specializing this equation to points A and B in Fig. 5D leads to (14) of the main text.

