## Supplemental Information for "Transcriptional competition biases the effects of second messengers in *Escherichia coli*"

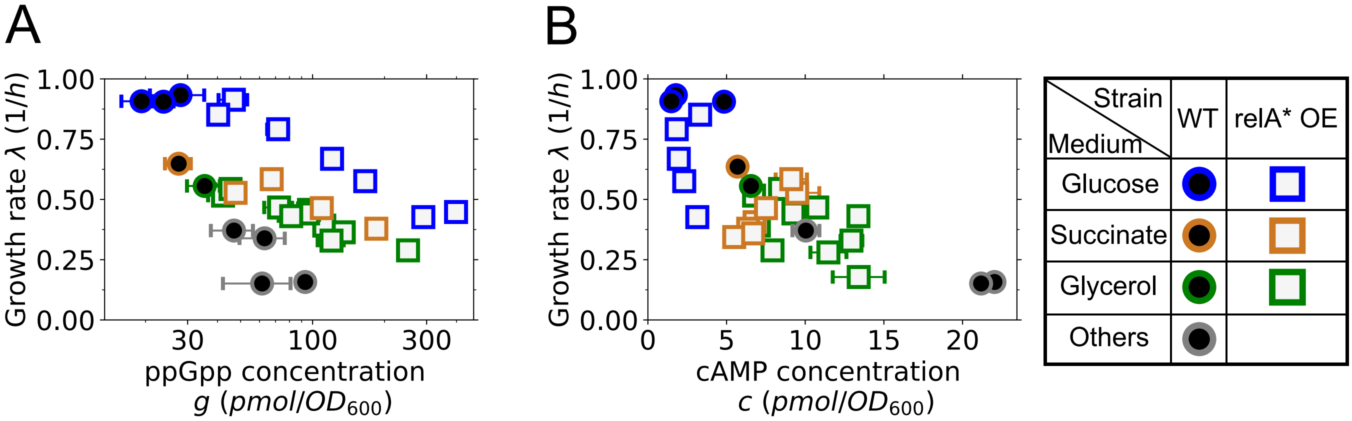


Figure S1: **Primary experimental data from this study: growth rates against ppGpp and cAMP concentration, related to Fig. 2, 3 and 4** A) The plot shows ppGpp concentration for cells growing in different carbon sources, both in the wild-type and in the RelA overexpression strain. Our results confirm the negative correlation between growth rate and ppGpp in the wild-type. We observe that RelA mutants deviate from the wild-type behaviour, with higher growth rates $\lambda$ at equivalent ppGpp, while still displaying a decrease. B) Same as plot A, for the second messenger cAMP. Our measurements show that cAMP concentration is negatively correlated with growth rate under catabolic limitation in the wild-type. In glucose, cAMP levels are unperturbed by RelA overexpression, while in succinate and glycerol the behavior with respect to growth rate is less defined, but cAMP levels show little variability compared to ppGpp (see also Fig. 3C).

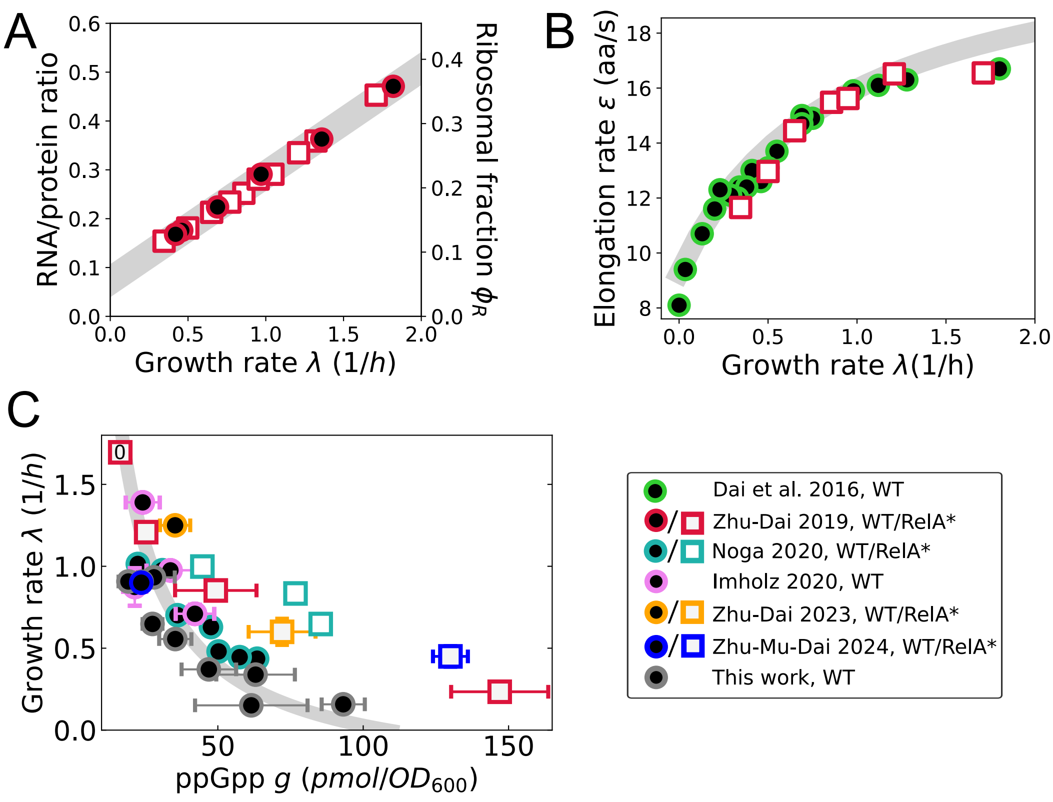


Figure S2: **Compilation of RelA overexpression data in *E.coli* NCM3722 from other studies, related to Fig. 2, 3, 4 and 5** A) The ribosomal growth law is conserved under RelA overexpression. This growth law was used in the main text to convert growth rate measurements into the corresponding ribosomal fractions. The grey line represents the best linear fit of wild-type data. Data from ref.[1] B) The wild-type elongation rate growth law $\epsilon(\lambda)$ (data from ref.[2]) is conserved under RelA overexpression (data from ref.[1]). The grey line represents (17), in agreement with ref.[2]. Note that our growth rate data under RelA overexpression fall within the $0.3 - 1\ h^{-1}$ range, where we can safely assume conservation of this growth law based on the available data. C) Deviation from the wild-type ribosome-ppGpp relationship under RelA overexpression. Wild-type data from this study and refs.[3,4] are interpolated as in Fig. 2B. RelA overexpression strain data are taken from refs.[1,4–6]. We denoted the point corresponding to the uninduced RelA overexpression strain in LB medium with a zero. This point, together with the wild-type data from refs[1,3,5,6], all fall along the interpolating line defined by our wild-type measurements.

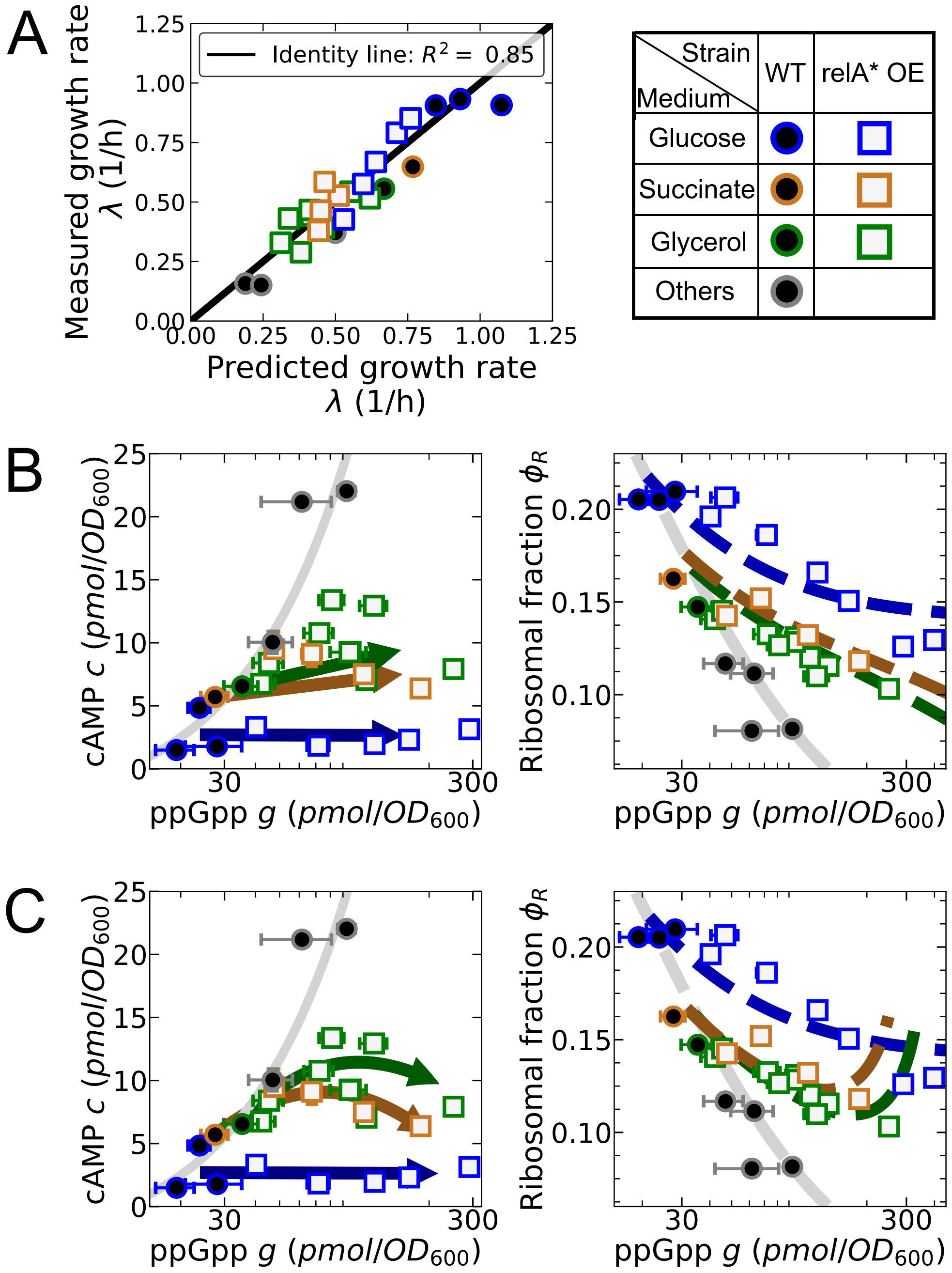


Figure S3: **Performance and visualization of the model, related to Fig. 3** A) The measured growth rate is plotted against the growth rate predicted by the model, with the parameters as in the main text, taking joint ppGpp and cAMP concentrations as inputs. All the points align along the identity line. B) While the goodness of fit is uniquely determined by the parameters, the visualization of the model's prediction for $\phi_R$ as a function of ppGpp depends on the fit chosen for cAMP as a function of ppGpp. If we force this fit to pass through wild-type points (or their average, for glucose), the outcome for $\phi_R$ is similar to the main text (Fig. 3A). C) Same as panel A, but here we fitted a quadratic cAMP-ppGpp relationship in succinate and glycerol. In this case, our model predicts minima for $\phi_R$ in these two carbon sources. The model still captures the behaviour of $\phi_R$ under extreme ppGpp overabundance.

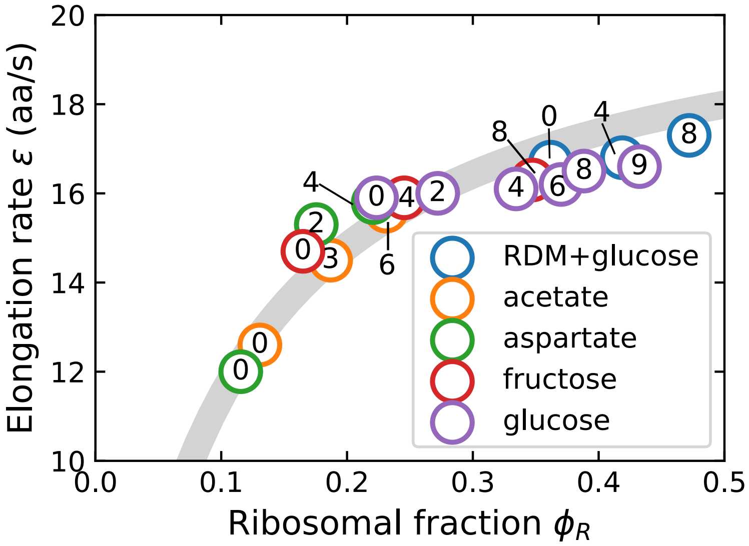


Figure S4: **The ribosome-elongation rate ($\phi_R - \epsilon$) relationship is conserved under chloramphenicol treatment. Related to Fig. 6**. Data from Dai et al.[2] for different nutrients, with varying chloramphenicol concentrations (annotated inside markers, $\mu M$). Chloramphenicol increases both translation elongation rate and amount of ribosomes according to the C-LIM law obtained by changing the carbon source without chloramphenicol ($0\ \mu M$ points). The gray line represents the best fit according to the C-LIM expression $\epsilon = \epsilon_{max}\frac{\phi_R}{\phi_R + k_M}$, with $\epsilon_{max} = 22$ aa/s and $k_M = 0.084$ as in ref.[2]. Ribosomal fraction values were obtain by conversion of measured RNA/protein ratios as in refs.[2,7].

### Note S1. Regulon-based proteome sectors

Related to Discussion and Fig S5.

To identify targets of ppGpp and cAMP, we leveraged the database RegulonDB[8] [1], which contains list of transcription factors with their target genes, and the mode of regulation (up- or down-regulation). When the mode of regulation was ambiguous, we excluded the gene from our analysis. We focused on how targets of cAMP-CRP and ppGpp-DksA are distributed across proteome sectors. For this analysis, we used the a more advanced model proposed by Mori et al.[10] where proteins are grouped into sectors based on their expression pattern (up- or down-regulation) under anabolic, catabolic, or translational limitation, as measured by proteomics. This method defines $2^3 = 8$ proteome sectors, among which we find the catabolic, the anabolic and the ribosomal [2] sectors.

Our analysis reveals that, while targets of repression by ppGpp are mainly ribosomal proteins, cAMP-CRP activation is not exclusive to the C sector, but widespread to sectors which responds very differently to catabolic limitation (Fig. S5B). We note in particular that cAMP activates components of the Q sector [3], and therefore its mass fraction might vary under RelA overexpression. The C sector comprises only 24% of the Crp-cAMP targets in the reference condition of the study by Mori et al. (glucose minimal medium). Even under conditions of severe catabolic limitation, it accounts for less than half of the targets (Fig. S5 A). In contrast, ppGpp targets are found mainly among ribosomal proteins or in the R sector defined as in Mori et al.[10] (Fig. S5B).

This observation prompted us to develop a more conservative and data-driven model. First, rather than considering the extended ribosomal sector, we consider only ribosomal proteins without affiliates. This is required because the ribosomal growth law under RelA overexpression[1] has been demonstrated to hold only for the RNA/protein ratio, which is proportional to the amount of ribosomal proteins. Although the assumption of proportionality between ribosomal and affiliated proteins under RelA overexpression, as made in the main text, is reasonable due to their coregulation[7], this relationship has yet to be validated specifically in the context of RelA overexpression. We can use the wild-type linear growth law for the restricted ribosomal sector (Fig. S5D, green line) as derived from the proteomics data in ref.[10] to convert our growth rate data into (restricted) ribosomal fractions, with the same method explained in the main text.

The second modification is that we replace the P sector with the cAMP regulon. More specifically, we consider only proteins that are (i) activated by cAMP-CRP according to RegulonDB and (ii) upregulated under catabolic limitation according to the proteomics data in Mori et al.[10]. This excludes genes for which cAMP cannot be considered as the primary regulator.

We note that this “cAMP-activated sector” is upregulated under RelA overexpression according to the data in ref.[6] (Fig. S5C), as predicted by our model through indirect effects (Fig. 3C). In the wild-type, its expression decreases with increasing growth rate under C-LIM (Fig. S5D, orange line, data from Mori et al.[10]). The total mass fraction of ribosomal proteins and cAMP activated sector is approximately constant and we use it as the new $\phi_{max} = 0.26$. We can use the same functional forms for $h_1(c)$ and $h_2(g)$ as in the main text to fit the data through our model of transcriptional competition. This model is still able to capture RelA overexpression data, with a change of parameters 1. In general, all our main conclusions remain valid, but this model can describe transcriptional competition only between ribosomal proteins and the cAMP regulon, rather than with the entire P sector.

[1] We repeated all the analysis with Ecocyc[9], with the same results.

[2] Note that with this method the ribosomal sector also contains some proteins which are not part of the ribosome nor ribosome-affiliated.

[3] We consider as belonging to Q all the sectors that do not respond specifically to one of the three limitations. More simply, we view Q as composed by C’, A’, S’, S, U.

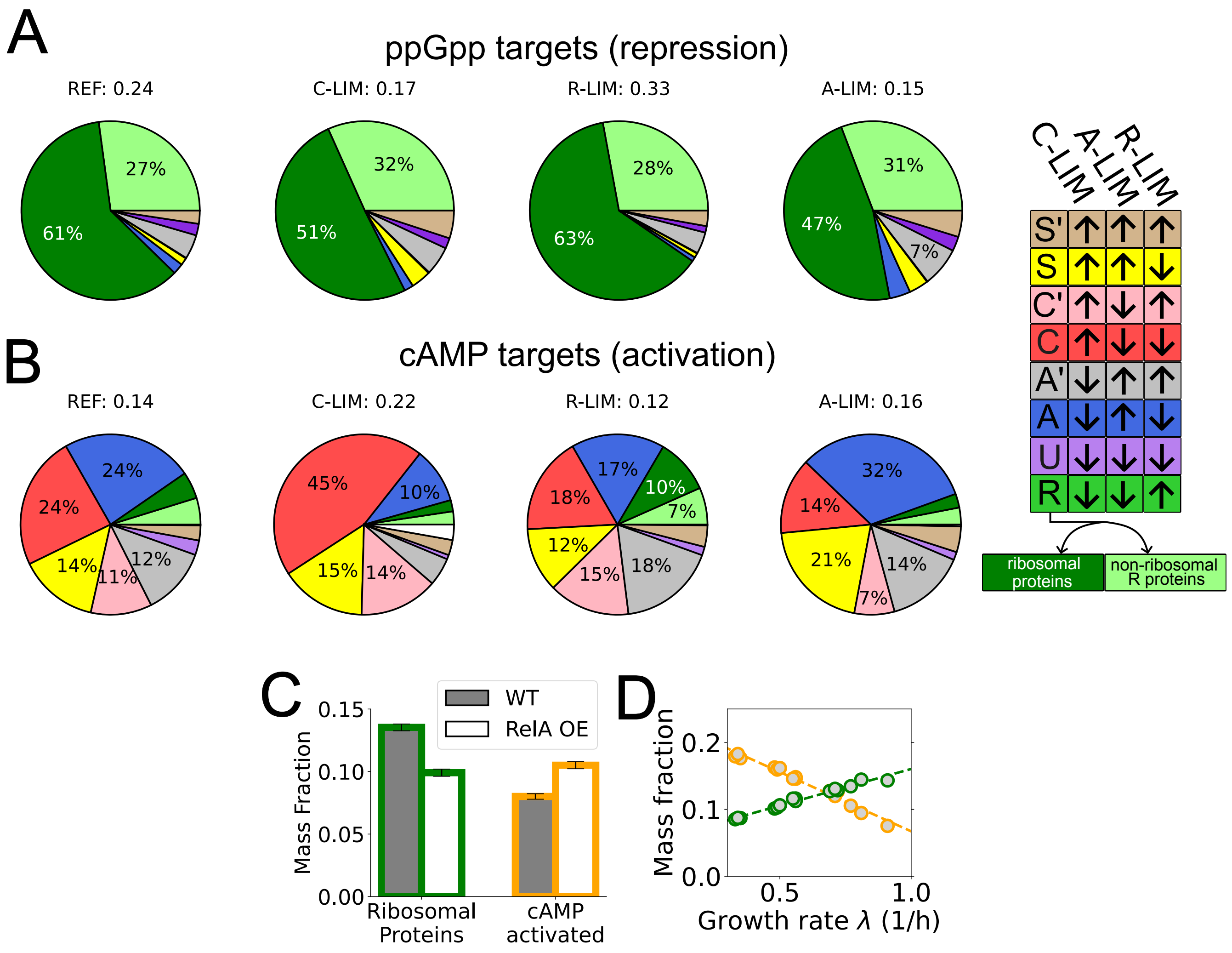


Figure S5: **cAMP-activated genes extend beyond the C sector and are upregulated under RelA overexpression according to the proteomics data in Zhu et al.[6]. Related to Fig 3 and Discussion.** **(A)** Our analysis of RegulonDB database[8] shows the distribution of ppGpp-repressed genes across proteome sectors as defined in Mori et al.[10] (see legend on the right). We distinguish between ribosomal proteins and other proteins in the R sector. **(B)** The same analysis for cAMP-activated genes reveals that these are scattered throughout different sectors. For A) and B), pie charts are weighted by protein abundance (data from Mori et al.[10]), which varies across conditions. We show results for the reference condition in Mori et al., as well as severe catabolic (C-LIM), translational (R-LIM), or anabolic (A-LIM) limitation. Numbers above pie charts indicate the total proteome fraction corresponding to cAMP or ppGpp targets in each given condition. **(C)** Data from Zhu et al.[6] show that RelA* overexpression in glucose minimal medium leads to repression of ribosomal proteins (dark green). At the same time, the "cAMP-activated sector" (orange) is upregulated. This sector is defined as the set of proteins activated by cAMP in Ecocyc and RegulonDB and upregulated under C-LIM in the wild-type. **(D)** In the refined variant of the model, ribosomal proteins (dark green) compete with the cAMP-activated sector (orange). The sum of the two sectors is approximately constant under nutrient limitation in the wild-type, with $\phi_{max} = 0.26$, data from Mori et al.[10]

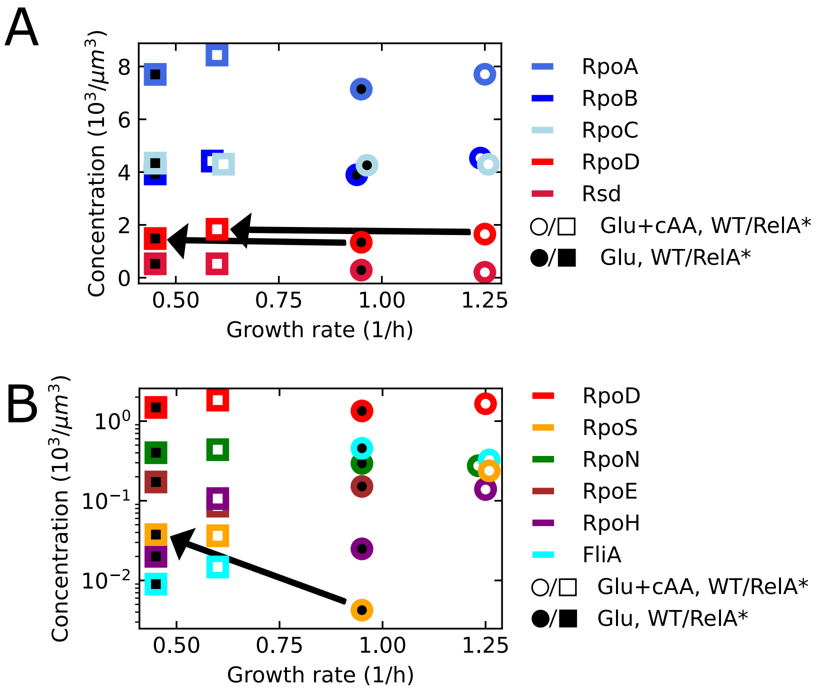


Figure S6: **RNAP and sigma factor landscape under RelA overexpression. Related to Fig. 3 and Discussion.** Data from ref.[5] (glucose+cAA, open markers) and ref.[6] (glucose, filled markers). Circles correspond to wild-type data, squares to RelA overexpression. A) RNAP core components (RpoABC) are in excess with respect to the housekeeping sigma factor RpoD. Free RpoD levels are reduced by its anti-sigma factor Rsd. RelA overexpression's effect on RpoD (arrows) and Rsd levels is negligible. B) Sigma factor landscape under RelA overexpression. The main change is an increase in RpoS abundance in glucose (as indicated by the arrow), while RpoD clearly remains the most abundant sigma factor (see the log scale). Note that rpoS can be knocked out without affecting the response to RelA overexpression[6], which we therefore expect to be independent of this sigma factor.

Table 1: **Table of parameters resulting from data fitting**

| **Fit parameters** | | | | | |
|---|---|---|---|---|---|
| Model | $\mu_1(-)$ | $\mu_2(pmol/OD_{600})$ | $\mu_3(pmol/OD_{600})$ | $\phi_{max}$ (imposed) | $R^2$ |
| No gene dose effects | 0.84 | 11.95 | 25.60 | 0.45 | 0.85 |
| With gene dose effects | 0.97 | 14.18 | 20.80 | 0.45 | 0.85 |
| Regulon-based model | 0.9 | 13.15 | 32.52 | 0.26 | 0.83 |
