## Supplementary material for "Transcriptional competition biases the effects of second messengers in *Escherichia coli*": Methods S1

*Detailed information regarding modeling*

### 1 Model for coarse-grained transcriptional competition

#### 1.1 Modeling transcription.

We rewrite and explain the dynamics of the concentration of a single species of transcripts as presented in the main text (3):

$$\frac{d[m_i]}{dt} = \alpha_i[G_i] - (\delta + \lambda)[m_i].$$

Here $\alpha_i$ is the initiation rate, and $[G_i]$ the gene concentration. As in refs.[1,2], we are considering an "initiation-limited" regime for transcription, where RNA polymerase traffic is negligible, and transcriptional elongation and termination do not create bottlenecks[3,4]. Thus, we neglect effects of RNAP pausing and traffic, which are expected to play a role in some circumstances, such as for rrn operons in fast growing *E.coli*[3]. For mRNA degradation we assume a sector-independent first-order kinetics with rate $\delta$ as in ref.[2]. The dilution term $-\lambda[m_i]$ accounts for the time variation in cellular volume. This term can be considered negligible with respect to mRNA degradation in *E.coli*[1,2].

In principle, the transcription initiation rate $\alpha_i$ depends on the availability of free RNA polymerases, including those non-specifically bound to DNA, as well as initiation factors and DNA properties, such as looping or the coiling state[5,6]. For simplicity, we consider initiation to be limited only by the diffusion of RNAP, and indicate the concentration of RNAP holoenzymes available to start transcription with $[N_{av}]$. We note that Balakrishnan et al.[1] showed that, in exponentially growing *E.coli*, $[N_{av}]$ is limited by the availability of free $\sigma^{70}$ factors, whose concentration is set by the anti-sigma factor Rsd. We show how alternative sigma factors can be incorporated into our framework in Section. 2.1.

Initiation of transcription by RNAP is a multi-step process that includes promoter binding, open complex formation, and promoter clearance[7]. A comprehensive expression for the initiation rate would be a Hill function, such as

$$\alpha_i = V_i \frac{[N_{av}]}{n_i + [N_{av}]}, \tag{S1.1}$$

where $V_i$ represents the clearance rate and corresponds to the maximal transcription rate[8]. The Michaelis-Menten factor reflects promoter occupancy, with the parameter $n_i$ modulating the promoter affinity for RNAP. Consistent with Balkrishnan *et al.*[1] and Calabrese *et al.*[2], we consider only the linear regime of this expression, a "weak promoter limit", which is a valid approximation for most mRNA promoters[5], and define the promoter strength

$$k_i = \frac{V_i}{n_i}, \tag{S1.2}$$

following ref.[3]. In this limit, the influence of RNAP abundance on proteome partitioning vanishes; and our framework remains applicable regardless of whether transcriptional regulation affects the clearance rate or promoter affinity. Transcript dynamics then becomes

$$\frac{d[m_i]}{dt} = k_i[N_{av}][G_i] - (\delta + \lambda)[m_i]\ . \tag{S1.3}$$

The next component of the model is coarse-graining genes into different sectors: transcriptome fractions are defined as $[m_I] := \sum_{j\in I}[m_j]$, and we obtain their evolution by summing (S1.3) for all the transcripts in a given sector,

$$\frac{d[m_I]}{dt} = \sum_{j\in I} k_j[G_j][N_{av}] - (\delta + \lambda)[m_I]\ . \tag{S1.4}$$

We define the coarse-grained sector strength

$$\beta_I := N_I \frac{\sum_{j\in I} k_j [G_j]}{\sum_{j\in I}[G_j]},$$

so that we can write

$$\frac{d[m_I]}{dt} = \beta_I [N_{av}] \frac{\sum_{j\in I}[G_j]}{N_I} - (\delta+\lambda)[m_I], \tag{S1.5}$$

and

$$\frac{d[m_I]}{dt} = \beta_I [N_{av}][G_I] - (\delta+\lambda)[m_i], \tag{S1.6}$$

which shows that $\beta_I$ can be interpreted as a coarse-grained promoter strength (compare with (S1.3)), provided that we define the typical gene concentration of sector I as

$$[G_I] = \frac{1}{N_I}\sum_{j\in I}[G_j]. \tag{S1.7}$$

Here $N_I$ is the total number of genes in sector I. We can also define the coarse-grained initiation rate $\alpha_I = \beta_I[N_{av}]$, which represents the initiation rate required to produce an equivalent expression (in terms of mRNAs) of sector I from a single gene, with the typical concentration $[G_I]$. This gives (5).

In the model presented in the main text, all the genes are assumed to be present at the same concentration, so that the typical concentration $[G_I]$ becomes sector-independent, $[G_I] = G$ for all I, and

$$\beta_I = \sum_{j\in I} k_j \; . \tag{S1.8}$$

Then the equilibrium of (S1.6) is given by

$$[m_I] = \frac{\beta_I[N_{av}][G]}{(\delta+\lambda)} \tag{S1.9}$$

and the total mRNA concentration is

$$[m_{TOT}] = \frac{\beta_{TOT}[N_{av}][G]}{(\delta+\lambda)} \; , \tag{S1.10}$$

with $\beta_{TOT} = \sum_I \beta_I$.

We rewrite the definition of transcriptome fractions given in the main text, $\chi_i := \frac{[m_i]}{[m]}$. Note that the same notation is typically used for ribosome allocation functions[9,10], which coincide with transcriptome fractions when ribosome binding is independent of the mRNA species. Their dynamics is

$$\frac{d\chi_I}{dt} = \frac{1}{[m]}\frac{d[m_I]}{dt} - \chi_i \frac{1}{[m]}\frac{d[m]}{dt}. \tag{S1.11}$$

From (S1.5) we get

$$\frac{d\chi_i}{dt} = \frac{\beta_I}{[m]}[G][N_{av}] - \chi_I \frac{\beta_{tot}}{[m]}. \tag{S1.12}$$

We rewrite the last equation as

$$\frac{d\chi_i}{dt} = \frac{\beta_{TOT}[G][N_{av}]}{[m]}\left(\frac{\beta_I}{\beta_{TOT}} - \chi_I\right) = \frac{1}{\tau_\chi}\left(\omega_I - \chi_I\right), \tag{S1.13}$$

with the definition

$$\omega_I := \frac{\beta_I}{\beta_{TOT}}. \tag{S1.14}$$

These functions, central to our framework, represent RNAP allocation on genes of different sectors, which coincides with ribosome allocation at steady-state (see (7)). They condense the effects of transcription factors, and are independent of growth rate or mRNA degradation rate. The relaxation timescale is defined as

$$\tau_\chi := \frac{[m]}{\beta_{TOT}[G][N_{av}]}, \tag{S1.15}$$

and, in general, it is not a constant. However, near the equilibrium point defined by (S1.10) we have

$$\tau_\chi \approx (\delta+\lambda)^{-1} \approx \delta^{-1}, \tag{S1.16}$$

consistent with ref[10].

#### 1.2 Modeling gene dosage effects

In *E. coli*, as in other fast-growing bacteria, DNA is replicated from a single origin, and gene copy numbers vary with the growth rate depending on their distance from the origin of replication[1,11,12]. Here, we provide a formal treatment of gene dosage effects, which were neglected in the main text due to their relatively small impact on gene expression compared to transcriptional regulation[1]. Gene concentrations can be written as relative to the concentrations of origins as

$$[G_i] = d(x_i,\lambda)[Ori]\ , \tag{S1.17}$$

where $d(x_j,\lambda) = e^{-\lambda C x_j}$ is the gene dosage relative to the origin of replication[1,12], The time interval $C$ is the C period, i.e., the time required for the two replication forks to traverse the chromosome from Ori to Ter. The product $\lambda C$ can be expressed as a linear function of the growth rate[1]. Fernandez-Coll[13] et al. showed that this dependency on the growth rate is conserved under ppGpp overabundance. When considering gene dosage effects, (S1.4) becomes

$$\frac{d[m_I]}{dt} = \sum_{j\in I} k_j\ d(x_j,\lambda)[Ori][N_{av}] - (\delta+\lambda)[m_j]\ . \tag{S1.18}$$

In STAR Methods, we have defined the "coarse-grained regulatory activities" (we follow the terminology of Balakrishnan *et al.*)

$$\theta_I := \sum_{j\in I} k_j \cdot d(x_j,\lambda),$$

so that we can express the coarse-grained transcript dynamics as

$$\frac{d[m_I]}{dt} = \theta_I[Ori][N_{av}] - (\delta+\lambda)[m_j]\ . \tag{S1.19}$$

To account for the contribution of gene dosage to proteome allocation, we adopt an approximate approach by assigning a coarse-grained positions $x_I$ to each sector, i.e. use

$$\theta_I \approx d(x_I,\lambda)\sum_{j\in I} k_j = d(x_I,\lambda)\beta_I\ , \tag{S1.20}$$

In this way we separate the contribution of transcriptional regulation (encoded by the dependency of $\beta_I$ on transcription factors such as ppGpp or cAMP) from the contribution of gene position (which depends on the growth rate through the function $d(x_I, \lambda)$. The equilibrium of (S1.19) is given by

$$[m_I] = \theta_I \frac{[Ori][N_{av}]}{\delta + \lambda} \tag{S1.21}$$

Following the same steps as in the previous section, we still find

$$\frac{d\chi_i}{dt} = \frac{1}{\tau_\chi} \left( \omega_I - \chi_I \right), \tag{S1.22}$$

with the definitions

$$\begin{aligned} \omega_I &:= \frac{\theta_I}{\theta_{TOT}} \\ \tau_\chi &:= \frac{[m]}{\theta_{TOT}[G][N_{av}]} \end{aligned} \tag{S1.23}$$

where

$$\theta_{TOT} := \sum_I \theta_I \ . \tag{S1.24}$$

In this variant, RNAP allocation functions $\omega_I$ are influenced by relative variations in gene dosage, but not by variations in Ori concentration, which affect all genes equally. Indeed

$$\omega_I = \frac{\beta_I \ d(x_I, \lambda)}{\sum_I \beta_I \ d(x_I, \lambda)}. \tag{S1.25}$$

When examining RNAP allocation, one can transition between models without gene dosage and with gene dosage effects with the substitution $\beta \to \theta$. We will address the discussion of gene dosage effects on our results in a later section.

#### 1.3 Modeling translation

As we did for transcription, we assume an initiation-limited regime for translation (see[1,2] for a similar treatment): we write the evolution of the concentration of proteins belonging to the I sector as

$$\frac{d[P_I]}{dt} = [m_I]\alpha_{TL,I}[R_{av}] - \eta_I[P_I] - \lambda[P_I] \tag{S1.26}$$

where $\alpha_{TL,I}$ is the translational initiation rate and $[R_{av}]$ is the concentration of free (available) ribosomes [1]. We include both the degradation term $\eta_I[P_I]$ and the dilution term $\lambda[P_I]$. As stated in the main text, we work under the assumption of absence of post-transcriptional regulation at a coarse level. Hence, the translation initiation rate $\alpha_{TL,I}$ and the degradation rate $\eta_I$ are sector-independent: $\alpha_{TL,I} = \alpha_{TL}$ and $\eta_I = \eta$. degradation effects vanish when considering the evolution of mass fractions, defined as $\phi_I := \frac{M_I}{M}$ (see (1)). We can transition between mass fractions and concentrations using the average protein lengths[14]:

$$\phi_I = \frac{[P_I]}{[P]}\frac{L_I}{L} \ , \tag{S1.27}$$

with $L_I$ and $L$ being the average protein length across the sector I and across the entire proteome, respectively. Mass fractions dynamics is given by

$$\frac{d\phi_I}{dt} = \frac{L_I}{L}\frac{T_I}{[P]}\alpha_{TL}[R_{av}] - \frac{L_I}{L}\frac{[P_I]}{[P]}\frac{[m]}{[P]}\alpha_{TL}[R_{av}], \tag{S1.28}$$

---
[1] More precisely, the concentration of available ribosomal subunits, assumed to be stoichiometrically balanced.

which we rewrite as

$$\frac{d\phi_I}{dt} = \frac{L_I}{L}\frac{T_I}{[m]}\frac{[m]}{[P]}\alpha_{TL}[R_{av}] - \frac{L_I}{L}\frac{[P_I]}{[P]}\frac{[m]}{[P]}\alpha_{TL}[R_{av}] \tag{S1.29}$$

and regroup as

$$\frac{d\phi_I}{dt} = \left(\frac{L_I}{L}\chi_I - \phi_I\right)\frac{\alpha_{TL}[R_{av}][m]}{[P]} \tag{S1.30}$$

The presence of the factor $\frac{L_I}{L}$ arises from our definition of transcriptome fractions; this factor vanishes in an approximate approach where proteins across different sectors are assumed to have the same average length.

Next, thanks to nonequilibrium traffic models[2], we can connect initiation and elongation:

$$\alpha_{TL}[R_{av}] = \frac{k_{TL}}{L}\frac{[R_a]}{[m]} \tag{S1.31}$$

where $k_{TL}$ is the speed of ribosomes on transcripts, in units of protein length per unit time, and $[R_a]$ is the concentration of actively translating ribosomes. We introduce the fraction of active ribosomes $f_a := \frac{[R_a]}{[R]}$ (for an alternative treatment, see ref.[2]).

We now introduce explicitly the ribosomal sector, which comprises ribosomal proteins and affiliates[9]. Since affiliates are in stoichiometric balance with ribosomal proteins, we can associate each ribosome with its extended protein mass[9] $m_{R,ext}$ and write the ribosome concentration as

$$[R] = \frac{1}{V}\frac{M_R}{m_{R,ext}}, \tag{S1.32}$$

while the total protein concentration can be expressed as

$$[P] = \frac{1}{V}\frac{M}{m}, \tag{S1.33}$$

with $m$ average protein mass. As a result,

$$\frac{d\phi_I}{dt} = \left(\frac{L_I}{L}\chi_I - \phi_I\right)\frac{k_{TL}}{L}\frac{m}{m_{R,ext}}f_a\phi_R \tag{S1.34}$$

We define the translational elongation rate:

$$\epsilon := \frac{k_{TL}}{L}\frac{m}{m_{R,ext}} = \frac{k_{TL}}{L_{R,ext}}, \tag{S1.35}$$

where $L_{R,ext}$ is the total length of an extended ribosome. Thus, $\epsilon$ can be interpreted as the inverse of the time needed for an extended ribosome to replicate itself [2]. We obtain the final expression

$$\frac{d\phi_I}{dt} = \left(\frac{L_I}{L}\chi_I - \phi_I\right)\epsilon f_a\phi_R \tag{S1.36}$$

As noted by[2], the length factor could be absorbed into the definition of promoter strengths: in fact, by retaining this factor, we find from the steady-state of the last equation

$$\phi_I = \frac{L_I}{L}\chi_I = \frac{L_I}{L}\omega_I = \frac{\beta_I L_I}{\beta_{tot}L}. \tag{S1.37}$$

[2]This equation converts the elongation rate from amino acids per second to elongation rate in $h^{-1}$, using the value $L_{R,ext} = 12307$ from ref.[15].

(compare with equation (7)). Here, we choose not to absorb this factor and instead carry it through our calculations in this document. In our code, we estimated $L_R$ from the average length of proteins in the "extended ribosomal sector" as defined in Scott et al.[9]. Considering that an extended ribosome contains $12307$ amino acids (aa)[15], we obtain $L_R = 166$ aa. To estimate $L_P = 365$ aa, we considered the list of genes assigned to the A and C sectors in Mori *et al.*.[16], calculating their average length weighted by abundance in the transcriptomic data of Balakrishnan et al.[1]. The length factor does not affect our results significantly (it only influences the fitted parameter $\mu_1$, see section 1.5).

#### 1.4 Sectors with constant size must be actively maintained at a fixed expression level

The extended form of (S1.37) for the Q sector reads

$$\phi_Q = \frac{L_Q}{L} \frac{\beta_Q}{\beta_R + \beta_P + \beta_Q} \tag{S1.38}$$

When the length factor is retained, the constraint $\phi_Q = const.$ yields

$$\beta_Q = \frac{\phi_Q}{\phi_{max}} \frac{\beta_R L_R + \beta_P L_P}{L_Q}, \tag{S1.39}$$

with the definition $\phi_{max} = 1 - \phi_Q$. A simple substitution into (S1.37) gives

$$\phi_R = \frac{\beta_R L_R}{\beta_R L_R + \beta_P L_P} \phi_{max} = \frac{1}{1 + \frac{\beta_P L_P}{\beta_R L_R}} \phi_{max} \tag{S1.40}$$

Note that the equation for $\phi_R$ is independent of $L_Q$. If protein length is considered the same for all sectors, one finds (8) of the main text.

#### 1.5 Fitting ribosomal fractions to the model of transcriptional competition

In the main text, we fit the model of transcriptional competition between P and R to our estimates for the ribosomal fraction $\phi_R(c, g)$. From (S1.40), we can see that all the mechanistic details of regarding transcriptional regulation are condensed into the "target function"

$$h(c, g) := \frac{\beta_P}{\beta_R} = \frac{L_R}{L_P} \left( \frac{\phi_R}{\phi_{max}} - 1 \right). \tag{S1.41}$$

which we try to replicate with our fit. This function completely determines $\phi_R$ from the hypothesis of a constant $\phi_{max} = 1 - \phi_Q$ (see (9)). Our fitting approach involves *separating* the effects of cAMP and ppGpp in $h(c, g)$ between numerator and denominator, which means that we seek two functions $h_1(c)$ and $h_2(g)$ such that

$$h(c, g) = \frac{h_1(c)}{h_2(g)}. \tag{S1.42}$$

Once their functional shapes have been chosen, $h_1(c)$ and $h_2(g)$ are completely determined by the fitting algorithm (Orthogonal Distance Regression). Then, as a first and basic guess, we assign

$$\beta_P(c) = h_1(c),\ \beta_2(g) = h_2(g) \tag{S1.43}$$

Choosing the functional shapes for these two functions completely determines the optimal fitting parameters ($\mu_1, \mu_2, \mu_3$ in the main text), the residuals and the goodness of the fit (Fig. S3A).

We note that, with the choice made in the main text, where the parameter $\mu_1$ has only a multiplicative role, any change in the ratio $\frac{L_R}{L_P}$ is completely absorbed by $\mu_1$. Thus, if the lengths are reparameterized, the effective concentrations $\mu_2$ $\mu_3$, as well as the goodness of fit, remain unaffected.

To visualize the model fit with continuity in the $\phi_R$-ppGpp plane, we used the following approach. For each carbon source, we fitted the cAMP-ppGpp relationship under RelA overexpression using a linear form, with $c = a \cdot g + b$, with different coefficients for each medium. We then computed $\phi_R$ along these fitted lines (denoted by the arrows in Fig. 3B) according to the model in equation 9, using parameters obtained from Orthogonal Distance Regression. The results are displayed as dashed lines in Fig. 3A. The same was done for wild-type points, yielding the grey dashed line in Fig. 3A. To ensure robustness in the visualization, we tried two other forms for the ppGpp-cAMP relationship. For instance, we forced intersection of wild-type points and assumed a medium-dependent linear (Fig. S3B) or quadratic (Fig. S3C) relationship under RelA overexpression. We stress that these choices affect only the visual representation of the ppGpp-$\phi_R$ fit, but not its quality.

#### 1.6 Fit including chromosomal position-dependent gene abundance

In the main text, we make the approximation that growth-rate variations in relative gene dosage are negligible when computing regulatory functions. This assumption is based on the narrow range of growth rates ($0.2 - 1$ h$^{-1}$) examined in this study. Nevertheless, we present the model variant including this factor and show that it does not impact our results significantly.

We recall that in our approximation regulatory activities are written as

$$\begin{aligned} \theta_R(g, \lambda) &= \beta_R(g) d(x_R, \lambda) \\ \theta_P(c, \lambda) &= \beta_P(c) d(x_P, \lambda). \end{aligned} \tag{S1.44}$$

To set values for the parameters $x_R$ and $x_P$, we computed sector-wide averages of gene positions weighted by mRNA abundance from data in[1], and verified that they show little dependence on the growth condition, producing $x_P = 0.6$, $x_R = 0.25$.

We retain the assumption $\phi_Q = const.$, which yields another version of (8):

$$\theta_Q = \frac{\phi_Q}{\phi_{max}} \left[ \beta_R(g) d(x_R, \lambda) \frac{L_R}{L_Q} + \beta_P(c) d(x_P, \lambda) \frac{L_P}{L_Q} \right] \tag{S1.45}$$

Therefore, the ribosomal fraction at steady state is, from (S1.23),

$$\phi_R(g, c, \lambda) = \frac{1}{1 + \frac{\beta_P(c) L_P}{\beta_R(g) L_R} D(\lambda)} \phi_{max} \tag{S1.46}$$

with $D(\lambda) := \frac{d(x_P.\lambda)}{d(x_R,\lambda)}$ accounting for different chromosomal position of genes in P and R.

In this variant, the target function is

$$h(c, g) := \frac{\beta_P(c)}{\beta_R(g)} = \frac{1}{D(\lambda)} \frac{L_R}{L_P} \left( \frac{\phi_{max}}{\phi_R} - 1 \right), \tag{S1.47}$$

and we can use it to repeat the same procedure exposed in the previous section. Our model is still able to capture RelA overexpression data, with a change of parameters. There is a significant decrease in the effective ppGpp concentration $\mu_3$ (Table 1). This occurs because regulation by gene dosage replaces part of the regulation by ppGpp needed to explain the data. It is important to note that the branching in Fig. 3A cannot be ascribed to gene dosage-dependent regulation (see Section 3.1).

### 2 Additional regulatory mechanisms

#### 2.1 Effects of alternative sigma factors and sigma factor competition

Here we discuss the robustness of our conclusions in the presence of alternative sigma factors. In the main text, we assume transcription to be limited by the housekeeping $\sigma^{70}$ (RpoD), as the model by Balakrishnan et al.[1] we build upon considers only this sigma factor to explain the physiology of wild-type, exponentially growing *E.coli*. This framework should remain valid even under RelA overexpression, provided ppGpp levels stay within the typical wild-type range (roughly below $100\,$pmol/OD$_{600}$, see Fig. 2B). Since the divergence in the $\phi_R$–ppGpp curve is already observable within this concentration range, our conclusion that ppGpp is not the sole determinant of ribosome abundance is not affected by neglecting alternative sigma factors in the model. In fact, the existence of alternative sigma factors reinforces the notion that the mapping between the concentration of a regulator (ppGpp) and its regulatees (ribosomal proteins) is not necessarily one-to-one, due to global effects.

However, alternative sigma factors may dominate physiology when ppGpp levels move outside the wild-type range, as many of them are upregulated by ppGpp. To examine this regime, we build on the sigma factor competition model of Mauri and Klumpp[17], which can be naturally integrated into our framework since it also assumes RNAP holoenzyme-limited transcription. In their model, a sigma factor $\sigma$ binds the RNAP core $E$ according to

$$[\sigma_{free}][E_{free}] \underset{K_U}{\overset{K_B}{\rightleftharpoons}} [\sigma E], \tag{S1.48}$$

where $E\sigma$ denotes the RNAP holoenzyme. The total (free and bound) concentration of a sigma factor is $[\sigma]$, and for each sigma factor one can define the equilibrium constant $K_{\sigma E} = k_U/k_B$. Given the complexity of sigma factor regulation, we introduce simplifying assumptions. First, as in the original model, we consider a single alternative sigma factor $\sigma_{ALT}$ competing with $\sigma^{70}$ for RNAP core binding, representing a lumped description of all alternative sigma factors. Second, we assume $\sigma_{ALT}$ does not overlap with the RpoD sigmulon and does not directly regulate genes in the P or R sectors, but instead activates a new proteome sector of mass fraction $\phi_{AL}$, and mRNA fraction $\chi_{ALT}$. We further assume the size of the Q sector, $\phi_Q$, is unaffected by sigma factor–dependent regulation. The initiation rate for gene $i$ is then written as

$$\alpha_i = k_{E\sigma,i}[G_i][E\sigma]. \tag{S1.49}$$

The only difference from Mauri and Klumpp is that we assume the weak promoter regime (as in Balakrishnan et al.[1]), so that $\alpha$ becomes a linear function of $[E\sigma]$. Incorporating ppGpp- and cAMP-dependent regulation of promoter affinities, and following the same steps leading to (S1.14), yields

$$\omega_R = \frac{\beta_R(g)[E\sigma^{70}]}{[\beta_R(g) + \beta_P(c)][E\sigma^{70}] + \beta_{ALT}[E\sigma^{ALT}]}\phi_{max} \tag{S1.50}$$

This equation implies that the *fractional* allocation of holoenzymes transcribing ribosomal genes, $\omega_R$, decreases if $\sigma^{ALT}$-dependent regulons are activated, all else being equal. This variable is central to our model, as at steady-state it determines the ribosomal fraction $\phi_R = \omega_R$. Note that this indirect effect does not necessarily arise from direct competition for the RNAP core. Rather, it may result from the normalization constraint on the fractional allocation of RNAP, which in turn determines transcription rates and proteome fractions. In other words, induction of $\sigma^{ALT}$-responsive genes can increase the number of their own transcripts without reducing the absolute number of $\sigma^{70}$-dependent transcripts. However, the *fraction* of $\sigma^{70}$-dependent (including ribosomal) mRNAs will decrease, leading to a corresponding reduction in their proteome fraction.

Note that this mechanism can also operate in the opposite direction: increasing or decreasing the activity of $\sigma^{70}$-dependent promoters can indirectly upregulate or downregulate $\sigma^{ALT}$-dependent genes, even in the absence of direct competition for the RNAP core. Indeed, the fractional allocation of RNAP to $\sigma^{ALT}$-responsive genes is

$$\omega_{ALT} = \frac{\beta_{ALT}[E\sigma^{ALT}]}{[\beta_R(g) + \beta_P(c)][E\sigma^{70}] + \beta_{ALT}[E\sigma^{ALT}]}\phi_{max}. \tag{S1.51}$$

Therefore, the fractional allocation of RNAP to $\sigma^{ALT}$-dependent genes can increase, for example, following a reduction in $\beta_R$ (due to elevated ppGpp) or a reduction in $\beta_P$ (due to decreased cAMP), while neglecting the direct effects of these second messengers on the concentration of $\sigma^{ALT}$ itself.

We can isolate the effect of competition for core RNAP by rewriting S1.50 as

$$\omega_R = \frac{\beta_R(g)}{[\beta_R(g) + \beta_P(c)] + \beta_{ALT}\frac{[E\sigma^{ALT}]}{[E\sigma^{70}]}}\phi_{max}, \tag{S1.52}$$

where all details of sigma factor competition are absorbed into the term $\frac{[E\sigma^{ALT}]}{[E\sigma^{70}]}$, which we leave implicit, as its dependence on global parameters is complex and analytical solutions are available only in special cases[17]. Indeed, competition between sigma factors also depends on sequestration during transcription and on the activity of anti-sigma factors, as discussed in Mauri and Klumpp[17], who noted that this effect can become significant when the total number of sigma factors approaches or exceeds the number of RNAP cores. Proteomics data[16] indicate that RNAP cores are present in large excess relative to sigma factors, and recent data by Zhu et al. suggest the same under RelA overexpression. Moreover, the only sigma factor that is significantly upregulated upon RelA overexpression in glucose is RpoS, which is not essential to explain proteomic reallocation as shown by knockout experiments[18].

Taken together, these observations suggest that while alternative sigma factors play an important role in stress adaptation and could, in principle, contribute to our experiments, they are not essential to explain the RelA overexpression phenotype. Although they can be incorporated into our framework, a detailed description of sigma factor competition beyond equation (S1.52) would require more quantitative knowledge of their regulation, including the effects of ppGpp and cAMP, which are not yet well defined.

Nonetheless, we examine here an interesting regime of competition originally analyzed by Mauri and Klumpp in the context of the stringent response. We consider the case where $\sigma^{ALT}$ has a lower affinity for the RNAP core than $\sigma^{70}$, while both binding affinities remain strong. This regime, which to some extent captures the competition between RpoD and RpoS[17], yields

$$\frac{[E\sigma^{ALT}]}{[E\sigma^{70}]} = \begin{cases} 0 & [E] < [\sigma^{70}] \\ \frac{[E]-[\sigma^{70}]}{[\sigma^{70}]} & [\sigma^{70}] < [E] < [\sigma^{70} + \sigma^{ALT}] \\ \frac{[\sigma^{ALT}]}{[\sigma^{70}]} & [E] > [\sigma^{70}] + [\sigma^{ALT}] \end{cases} \tag{S1.53}$$

Strong competition occurs in the first two regimes. When the core deficit is severe ($[E] < [\sigma^{70}]$), all RNAP cores are bound to the strong housekeeping sigma factor, with no binding to the alternative one. In the intermediate regime ($[\sigma^{70}] < [E] < [\sigma^{70}] + [\sigma^{ALT}]$), the number of cores is sufficient to bind all $\sigma^{70}$ molecules but not all $\sigma^{ALT}$, so the amount of bound $[E\sigma^{ALT}]$ is determined by the cores remaining after $\sigma^{70}$ binding. When competition is weak, that is, when enough cores are available to bind all sigma factors, the ratio between the bound holoenzymes reflects the ratio of their total abundances. Notably, in this simplified framework, the weak-competition regime is the only one in which $\sigma^{ALT}$ explicitly appears in (S1.52). This implies that titration of a sigma factor weaker than the housekeeping one can substantially influence gene expression only under conditions of RNAP core excess.

#### 2.2 Common regulatory factors only affect the total regulatory activity

Consider again equations S1.42. Instead of S1.43, which we rewrite below

$$\beta'_P(c) = h_1(c),\ \beta'_R(g) = h_2(g)$$

we can also write, in absence of ulterior constraints,

$$\beta'_P(c) = \rho \cdot h_1(c),\ \beta'_R(g) = \rho \cdot h_2(g) \qquad \text{(S1.54)}$$

where $\rho$ is a completely free parameter, which could in principle depend on the single data point under evaluation. This means that in our fitting approach is unable to identify regulatory factors or mechanisms that affect both sectors in a similar way (e.g., a hypothetical transcription factor repressing both metabolism and ribosomes). Note that two choices (S1.43) and (S1.54) differ when the total promoter strength is computed:

$$\begin{aligned}
\beta_{tot} &= \frac{1}{\phi_{max}}\left[h_1(c) + h_2(g)\right] \\
\beta'_{tot} &= \frac{1}{\phi_{max}}\rho\left[h_1(c) + h_2(g)\right],
\end{aligned} \qquad \text{(S1.55)}$$

and therefore produce two different total regulatory activities:

$$\begin{aligned}
\theta_{TOT} &= \frac{1}{\phi_{max}}\left[h_1(c)d(\lambda, x_P) + h_2(g)d(\lambda, x_R)\right] \\
\theta'_{TOT} &= \frac{1}{\phi_{max}}\rho\left[h_1(c) \cdot d(\lambda, x_P) + h_2(g) \cdot d(\lambda, x_R)\right],
\end{aligned} \qquad \text{(S1.56)}$$

with

$$\mathrm{FC}(\theta_{TOT}) = \mathrm{FC}(\rho) \cdot \mathrm{FC}\{[h_1(c)d(\lambda, x_P) + h_2(g)d(\lambda, x_R)]\}. \qquad \text{(S1.57)}$$

hus, if such additional regulation exists, our model still reproduces the data; however, the sum $h_1(c) + h_2(g)$ does not represent the total promoter strength or the total regulatory activity.

#### 2.3 Common regulators and non-orthogonal effects

The $\rho$ factor accounts for the influence of transcription factors affecting both sectors, or transcription effects influencing e.g. supercoiling/DNA compaction. In addition to these, non-orthogonal effects by ppGpp on the P sector or by cAMP on the R sector could contribute to $\rho$. These effects cannot be direct, because we did not find a significant overlap between cAMP-activated regulon and ppGpp-repressed regulon (Fig. S5A,B). However, ppGpp has been reported to interfere with Crp[19]. In principle, this implies that the same cAMP concentration can correspond to different levels of activation of the catabolic sector if ppGpp levels are different. Variations in Crp levels are not explicitly incorporated into our framework, as we consider cAMP to be the effective regulator and do not explicitly model Crp. Nevertheless, to illustrate this scenario, we approximate and represent the strength of the catabolic sector as a function of both cAMP and ppGpp.

$$\beta_P = h_1(c) \cdot \zeta(g) \qquad \text{(S1.58)}$$

with $\zeta(g)$ decreasing with increasing ppGpp, representing ppGpp effects on CRP activity. Therefore, the fitted function $h_2(g)$ does not correspond to the "true", mechanistic, $\beta_R$, because the separation described in section 1.5 produces

$$h_2(g) = \frac{\beta_R(g)}{\zeta(g)}, \qquad \text{(S1.59)}$$

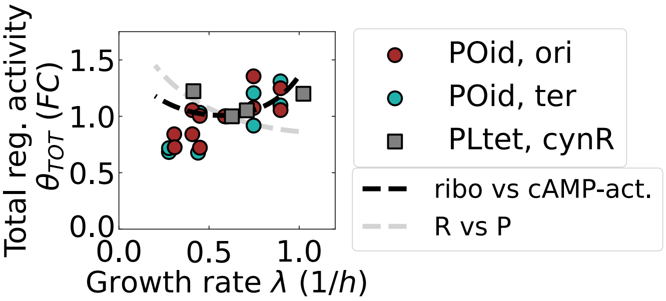

Figure S1.1: **Estimates for the total regulatory activity obtained from data in Balakrishnan et al.[1] and Zhang et al.[20]**. These estimates are based on the expression of three constitutive promoters inserted in three different chromosomal loci, plotted against the prediction of our models using the parameters obtained through fitting. Estimates and prediction are normalized for $\lambda = 0.6\ h^{-1}$, the midpoint of the interval of interest. The refined model (ribosomal proteins vs cAMP-activated sector) better captures the data, with respect to the R vs P model of the main text.

(still consistent with $h(c,g) := \frac{\beta_P}{\beta_R} = \frac{h_1(c)}{h_2(g)}$). In this case, $h_2(g)$ overestimates the repressive effect by ppGpp on the R sector. The total regulatory activity is

$$\beta'_{tot} = \zeta(c)\,[h_1(c) + h_2(g)] \tag{S1.60}$$

with $\zeta(c)$ being a repression at slow growth in the wild-type.

#### 2.4 Estimating fold-changes in the total regulatory activity.

To assess whether our data support the presence of additional regulation, we compare the total regulatory activity obtained with $\rho = 1$ in the wild-type to the values estimated by Balakrishnan et al.[1]. Below, we outline their estimation approach.

Consider a constitutive promoter, with its associated promoter strength $k_0$ constant across condition. At steady state this promoter drives the expression of a mass fraction

$$\phi_0 = \frac{L_0}{L}\frac{k_0 d(x_0,\lambda)}{\theta_{TOT}}, \tag{S1.61}$$

where $g_0$ is the relative gene concentration of the promoter. If the chromosomal position of the constitutive promoter is known, it is possible to estimate fold-changes in the denominator $\theta_{TOT}$ across conditions through the variations in the expression $\phi_0$:

$$\text{FC}(\theta_{TOT}) = \frac{\text{FC}(d(x_0,\lambda))}{\text{FC}(\phi_0))}. \tag{S1.62}$$

Balakrishnan et al. show that these estimates are consistent with respect to the insertion of the construct in different chromosomal loci. The resulting total regulatory activity is increasing with the growth rate (Fig. S1.1). We note that that the behaviour of constitutive promoters at slow growth can vary across different studies and so does the estimate for the total regulatory activity; we report for example estimates obtained from the expression of a Pltet promoter in Zhang et al.[20], where the total regulatory activity has a slightly different trend at slow growth.

If $\rho = 1$, our model predicts, through a combination of (S1.24) and (S1.45):

$$\theta_{TOT} = \beta_R d(x_R,\lambda) + \beta_P d(x_P,\lambda) + \frac{\phi_Q}{\phi_{max}}\left[\beta_R d(x_R,\lambda)\frac{L_R}{L_Q} + \beta_P d(x_P,\lambda)\frac{L_P}{L_Q}\right], \tag{S1.63}$$

or equivalently through (S1.46)

$$\theta_{TOT} = \frac{\beta_P \cdot d(x_P, \lambda)}{\phi_P(\lambda)} \left[ \frac{\phi_R(\lambda)}{L_R} + \frac{\phi_P(\lambda)}{L_P} + \frac{\phi_Q}{L_Q} \right], \tag{S1.64}$$

which connects total regulatory activity and promoter strength for a single sector (here the P sector). The main text model (including the A sector) predicts a decrease in the total regulatory activity with the growth rate (Fig. S1.1, grey line), when the estimates point towards a moderate increase for $\lambda \approx 1\, h^{-1}$. Restricting the competition to cAMP-activated targets,(thus excluding the anabolic sector), reconciles our results with the expected behaviour of $\theta_{TOT}$. We conclude that assuming $\rho = 1$ (absence of non-orthogonal cAMP-ppGpp effects) is an adequate approximation for our model.

### 3 Models without transcriptional competition with cAMP targets are unable to explain the data

#### 3.1 RelA overexpression data cannot be explained by gene dosage effects.

To exclude the possibility that the branching in 3A is determined by gene dosage effects, consider the wild-type and RelA overexpression mutants growing at the same rate $\lambda$, and hence with the same $\phi_R$ (Fig. S1.2B). The two situations correspond to different ppGpp concentrations, but the same factor $D(\lambda)$, because growth rate dependence of the ori/ter ratio remains unchanged under ppGpp overabundance[13]. Recalling (S1.46)

$$\phi_R(g, c, \lambda) = \frac{1}{1 + \frac{\beta_P L_P}{\beta_R L_R} D(\lambda)} \phi_{max},$$

the only way for both conditions to maintain the same $\phi_R$ despite different ppGpp levels (which affect $\beta_R$) is through a compensatory change in $\beta_P$. This implies that the P sector is not passive but actively competes with ribosomal proteins, consistent our theory.

#### 3.2 RelA overexpression data cannot be explained by ppGpp-driven activation of the P sector

Previous works suggest that anabolic and catabolic genes are activated by ppGpp[21]. We show here that a model where the P sector is activated by ppGpp, but cAMP effects are neglected, cannot explain the data. In this scenario, the P sector strength is an increasing function of $g$, $\beta_P = \beta_P(g)$, and equation (7) can be written as

$$\phi_R = \frac{\beta_R(g)}{\beta_R(g) + \beta_P(g)} \phi_{max} = \phi_R(g); \tag{S1.65}$$

therefore, $\phi_R$ becomes a function of ppGpp alone. According to this model, ppGpp titration by RelA overexpression should preserve the wild-type $\phi_R - g$ relationship, contradicting our data (Fig. 3A). Indeed, in this model, the activation of the P sector $\beta_P$ at a given ppGpp concentration should be the same in RelA overexpression mutants and in the wild-type, imposing the same degree of indirect repression on $\phi_R$.

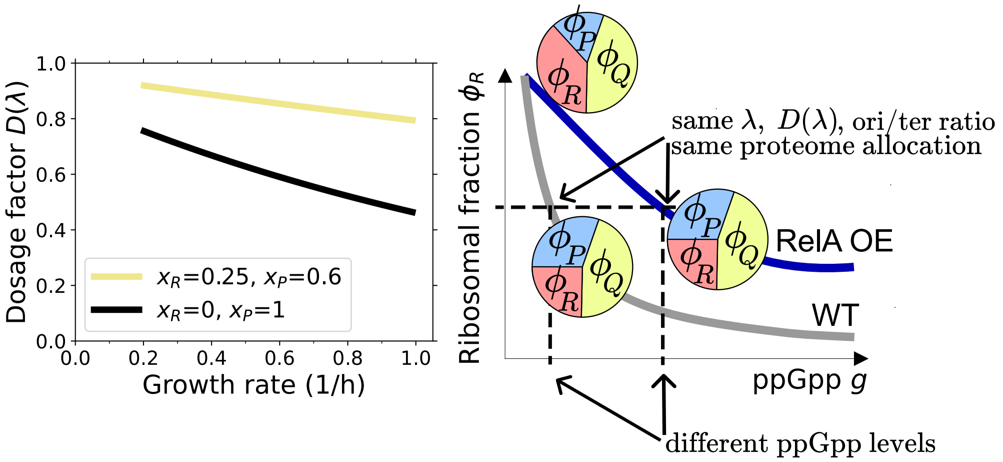

Figure S1.2: **The divergence in the ppGpp-$\phi_R$ relationship is not caused by gene dosage effects.** A) Dependence of the dosage factor on the growth rate, for $x_P$ and $x_R$ computed from transcriptomics data in ref.[1] (yellow line). The factor corresponding to the maximal distance (ori-ter) is also plotted for comparison. B) Gene dosage effect cannot explain the branching in $\phi_R - g$. At equivalent ribosomal fractions, the dosage factor $D(\lambda)$ is the same in wild-types and RelA overexpression mutants, but ppGpp concentration is different. Lines represent trends in data points in Fig. 3A, taking glucose as a reference.

However, we note that a model including both ppGpp and cAMP activation of the P sector is possible and would be able to capture the divergence: in this case $\beta_P = \beta_P(c, g)$ and

$$\phi_R = \frac{\beta_R(g)}{\beta_R(g) + \beta_P(c, g)} \phi_{max} = \phi_R(c, g), \tag{S1.66}$$

with ribosome abundance now depending on cAMP. Fully specifying such a model requires detailed knowledge of ppGpp and cAMP regulation across catabolic and anabolic genes, e.g., the fraction of genes responsive to both signals, the quantitative strength of their regulation, and the presence of possible doubly regulated targets. Given these complexities, we refrain from expanding our work in this direction, as the minimal model with cAMP activation alone is already sufficient to account for both published observations[22,23] and our new data.

#### 3.3 RelA overexpression data cannot be explained by an aberrant Q sector if P is passive.

As discussed in the main text, in order to account for the branching in Fig 3A, rather than introducing transcriptional regulation for the P sector, one can consider relaxing the assumption $\phi_Q = const.$ under RelA overexpression. This is the same as breaking the relation (8), which states that the strength $\beta_Q$ is adjusted to compensate for variations in $\beta_R$ and $\beta_P$. To avoid confusion, in this section we denote with $\phi_Q^{WT}$ and $\phi_{max}^{WT}$ the wild-type mass fraction of the housekeeping sector and its complementary fraction.

We first study a model variant where the P sector is constitutive-like, with a constant $\beta_P$ (therefore this variant does not contain cAMP regulation), and $\beta_Q$ remains unperturbed under RelA overexpression (therefore the Q sector size does not adjust to the variation in total promoter strength/regulatory activity). We show that this model is unable to explain the branching in Fig. 3A.

Consider the wild-type in a fast-growth condition (e.g., glucose medium), and let $g_f$ be the ppGpp concentration. Let $g_s > g_f$ be the ppGpp concentration in a condition of slower growth.

This concentration can also be obtained by inducing RelA from the fast-growth condition (Fig. S1.3).

Denoting with $\phi^*_R$ the ribosomal fraction under RelA overexpression, we have

$$\phi^*_R(g_s) = \frac{\beta_R(g_s)}{\beta_P + \beta_R(g_s) + \beta_Q(g_f)} = \frac{\beta_R(g_s)}{\beta_P + \beta_R(g_s) + \frac{\phi_Q^{WT}}{\phi_{max}^{WT}}[\beta_P + \beta_R(g_f)]}, \qquad \text{(S1.67)}$$

where we used the assumption that $\beta_Q$ remains unperturbed and expressed it with equation (8). We compare this fraction with the wild-type at the same ppGpp concentration $g_s$:

$$\phi_R(g_s) = \frac{\beta_R(g_s)}{\beta_P + \beta_R(g_s)}\phi_{max}^{WT} = \frac{\beta_R(g_s)}{\beta_P + \beta_R(g_s) + \frac{\phi_Q^{WT}}{\phi_{max}^{WT}}[\beta_P + \beta_R(g_s)]}. \qquad \text{(S1.68)}$$

We can rewrite $\phi^*_R(g_s)$ as

$$\phi^*_R(g_s) = \frac{\beta_R(g_s)}{\beta_P + \beta_R(g_s) + \frac{\phi_Q^{WT}}{\phi_{max}^{WT}}(\beta_P + \beta_R(g_s)) + \frac{\phi_Q^{WT}}{\phi_{max}^{WT}}[\beta_R(g_f) - \beta_R(g_s)]}. \qquad \text{(S1.69)}$$

Comparing the last two equations, under RelA overexpression $g_s > g_f$, thus $\beta_R(g_s) < \beta_R(g_f)$ and

$$\phi^*_R(g_s) < \phi_R(g_s). \qquad \text{(S1.70)}$$

Therefore, according to this model variant, we would expect a **stronger** repressive effect by ppGpp in RelA mutants. This contradicts experimental observations and we reject this scenario.

Our mathematical observations can be rationalised as follows. In the wild-type, if P is passive, constancy of Q sector size requires upregulation with decreasing ppGpp to counteract the de-repression of the R sector. This means that $\beta_Q$ increases with decreasing ppGpp. If under RelA overexpression Q remains activated at the same level, but competes with a repressed R sector, this results in a stronger repression of R with respect to the wild-type, as shown in Fig. S1.3.

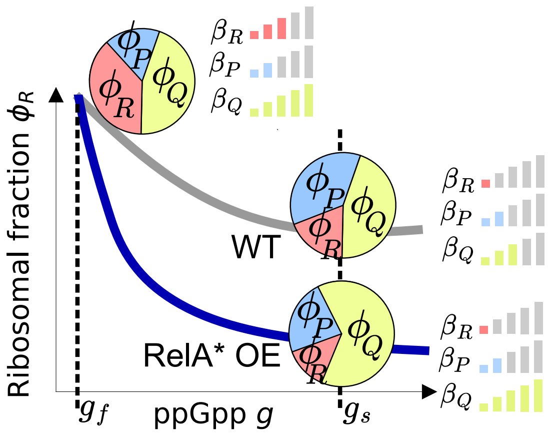

Figure S1.3: **Aberrant regulation of the Q sector cannot explain our data if the P sector is treated as constitutive.** Here we consider a model where P is constitutive, and the Q sector promoter strength remains unaltered under RelA overexpression, while still following (8) in the wild-type. The mass fraction $\phi_Q$ varies under overexpression and, at equivalent ppGpp concentration, ribosomes are *more* repressed in RelA OE mutants compared to the wild-type, in contrast with our data (compare with Fig. 3)

#### 3.4 RelA overexpression data cannot be explained by competition with the Q sector only

In the main text, the assumption $\phi_Q = const.$ implies that, under overexpression, the proteomic fraction “lost” by ribosomes is redirected towards the P sector. To strengthen the robustness of our findings, we can explore a model where this space is instead filled by the Q sector by imposing $\phi_P = const.$ under overexpression. This variant is inspired by the findings of Zhu et al.[18], who proposed that, in a regime of extreme ppGpp overabundance, downregulation of ribosomal genes is mainly due to activation of a stres response, rather than metabolism. We demonstrate that this scenario still requires competition between ribosomal proteins and cAMP regulon in the wild-type to explain the data.

We use a proof by contradiction: assuming that the P sector is passive ($\beta_P = const.$), while its mass fraction $\phi_P$ remains constant under RelA* overexpression, we demonstrate via the scheme in Fig. S1.4 that these assumptions are mutually inconsistent. Point X and Y represent the wild-type in a fast or slow growth condition, respectively. The R sector is repressed by ppGpp, so $\beta_R(X) > \beta_R(Y)$. If P were passive, this would imply

$$\beta_Q(X) > \beta_Q(Y) \tag{S1.71}$$

due to (8), which holds for wild-type points. Consider now point Z, which corresponds to RelA mutants at the same ppGpp concentration of point Y (slow growth condition in the wild-type), which gives the same $\beta_R$. Since $\beta_P$ is assumed to be constant, but $\phi_R(Z) > \phi_R(Y)$ (i.e., we observe branching under RelA* overexpression), we must have

$$\beta_Q(Z) < \beta_Q(Y)\ , \tag{S1.72}$$

which results from $\phi_R = \frac{\beta_R}{\beta_R+\beta_Q+\beta_P}$ (equilibrium equation for the ribosomal fraction). In simple terms, (S1.72) means that, at equivalent ppGpp levels, branching is due a stronger activation of the Q sector in RelA* mutants.

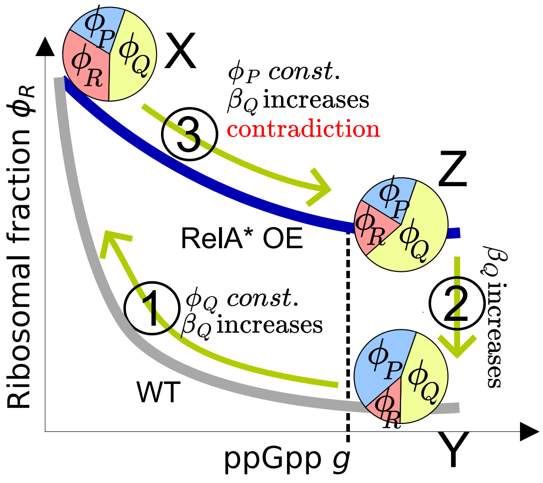

Figure S1.4: **Competition with the Q sector only does not explain RelA overexpression data if the P sector is treated as constitutive.** This figure illustrates the chain of relationships that leads to contradiction.

Finally, considering the constraint $\phi_P = const.$ under overexpression (X→Z), we would have

$$\frac{\beta_P}{\beta_P + \beta_R(X) + \beta_Q(X)} = \frac{\beta_P}{\beta_P + \beta_R(Z) + \beta_Q(Z)} \tag{S1.73}$$

(equilibrium equation for $\phi_P$), which implies

$$\beta_Q(X) < \beta_Q(Z), \tag{S1.74}$$

because $\beta_R(X) > \beta_R(Z)$ due to ppGpp's effect. Note that (S1.71), (S1.72) and (S1.74) are incompatible. Therefore, the assumption $\beta_P = const.$ is unable to explain the data in this scenario, which requires active regulation of the P sector, consistent with activation by cAMP, $\beta_P = \beta_P(cAMP)$, as in our theory.

### 4 Estimating the fold change in RelA* synthase activity

In this section, we estimate the relative changes in ppGpp synthase activity of overexpressed RelA* using our growth rate and ppGpp data, following the model by Wu *et al.*[24]. In their framework, ppGpp synthesis by endogenous RelA is given by

$$a = k_a[R^a_{dwell}] \tag{S1.75}$$

where $[R^a_{dwell}]$ represents the concentration of active "dwelling" ribosomes, and $k_a$ is a proportionality constant. Similarly, ppGpp degradation follows first-order kinetics, with first-order rate coefficient

$$b = k_b[R^a_{trans}] \tag{S1.76}$$

where $[R^a_{trans}]$ is the concentration of translocating ribosomes, and $k_b$ as the proportionality constant. While Wu *et al.* express these rates in terms of ribosome numbers, we use concentrations because we directly measured ppGpp concentrations in our experiments. This approach is equivalent to theirs, assuming ppGpp dilution by volume growth is negligible. Thus, in wild-type cells we have

$$\frac{dg}{dt} = k_a[R^a_{dwell}] - k_b[R^a_{trans}]g \ , \tag{S1.77}$$

and the steady-state ppGpp concentration is given by

$$g_{eq} = g_0 \frac{[R^a_{dwell}]}{[R^a_{trans}]}, \tag{S1.78}$$

where $g_0 = \frac{k_a}{k_b}$. Wu *et al.* introduce the characteristic "dwelling time" ($\tau_{dwell}$), the waiting period for cognate tRNA binding, and the "translocation time" ($\tau_{trans}$), which includes peptidyl transfer and translocation. The translocation time corresponds to the inverse of the maximal elongation rate, $\tau^{-1}_{trans} = \epsilon_{max}$. Since protein elongation involves both dwelling and translocation phases, the measured elongation rate $\epsilon$ follows

$$\epsilon^{-1} = \tau_{dwell} + \tau_{trans}. \tag{S1.79}$$

By applying ribosome flux balance, i.e.

$$[R_{dwell}]\tau^{-1}_{dwell} = [R_{trans}]\tau^{-1}_{trans} \tag{S1.80}$$

Wu *et al.* obtain for the wild-type

$$\frac{g}{g_0} = \frac{\tau_{dwell}}{\tau_{trans}} = \frac{\epsilon_{max}}{\epsilon} - 1 \ , \tag{S1.81}$$

as shown in Fig. 4A.

To incorporate the effect of RelA* overexpression, we modify (S1.77) by introducing an additional ppGpp synthesis term, $k_*$, proportional to the concentration of the constitutively active[25,26] RelA* expressed from the pTet promoter, as in (12) of the main text:

$$\frac{dg}{dt} = k_a[R^a_{dwell}] + k_* - k_b[R^a_{trans}]g. \tag{S1.82}$$

Using ribosome flux balance, this can be written as

$$\frac{dg}{dt} = [R^a]\frac{k_a\tau_{dwell} - k_b\tau_{trans}\frac{g}{g_0}}{\tau_{dwell} + \tau_{trans}} + \frac{1}{k_a}k_*. \tag{S1.83}$$

Rewriting in terms of elongation rates, thanks to (S1.79):

$$\frac{dg}{dt} = \frac{[R^a]\epsilon}{\epsilon_{max}}\left[\left(\frac{\epsilon_{max}}{\epsilon} - 1\right)\frac{g}{g_0}\right] + \frac{1}{k_a}k_*. \tag{S1.84}$$

At steady state $\frac{dg}{dt} = 0$, we solve for $k_*$:

$$k_* = k_a\frac{[R^a]\epsilon}{\epsilon_{max}}\left[\frac{g}{g_0} - \left(\frac{\epsilon_{max}}{\epsilon} - 1\right)\right] \propto \lambda\left[\frac{g}{g_0} - \left(\frac{\epsilon_{max}}{\epsilon} - 1\right)\right]. \tag{S1.85}$$

In the above equation, we used the proportionality

$$[R^a]\epsilon \propto \lambda \tag{S1.86}$$

which follows from the relation between mass fraction of active ribosomes, elongation rate and growth rate

$$\phi^a_R\epsilon = \lambda. \tag{S1.87}$$

The last two equations are equivalent because mass fractions and concentrations are interchangeable[14], assuming negligible changes in total protein concentration when RelA* is overexpressed. The final version of (S1.85) allows us to estimate fold changes in $k_*$ using growth rates, elongation rates, and ppGpp concentrations. Intuitively, (S1.85) means that RelA* activity corresponds to the vertical difference between mutant and wild-type data in Fig. 4B, scaled by the mutant growth rate. Consistent with our expectations, Fig. 4 demonstrates that the activity increases with the relative concentration of doxycycline, which we use as a broad proxy for the level of RelA* induction.

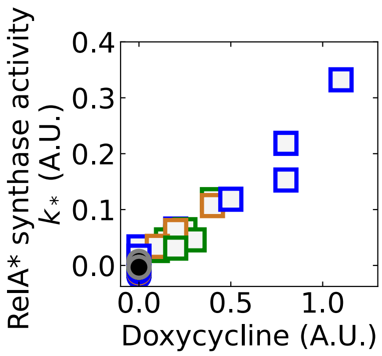

Figure S1.5: **RelA* activity as estimated from our data through** (S1.85) **plotted against the inducer (doxycycline) level.** Legend as in Fig 3. Wild-type points collapse at the origin, while $k_*$ is positively correlated with doxycycline for all carbon sources in RelA overexpression mutants.

### 5 Growth limitations and transcriptional competition

#### 5.1 Anabolic limitation

In the main text, we divide the metabolic P sector into its anabolic (A) and catabolic (C) components. We explicitly associate these two sectors with the strengths $\beta_A$ and $\beta_C$, with $\beta_P = \beta_A + \beta_C$. Kochanowski et al.[23] suggested that the A sector can be considered constitutive and passively regulated. This is based on the observation that anabolic enzymes and constitutive proteins show similar expression trends under cAMP titration, indicating that $\beta_A$ is poorly influenced by this transcription factor. The same study confirms that catabolic enzymes can be seen as activated by cAMP, with $\beta_C = \beta_C(c)$. Therefore, in our model the P sector strength $\beta_P$ is responsive to cAMP as well. We note that with this configuration, while the total metabolic fraction $\phi_P$ is indirectly influenced by ppGpp (due to competition with $\phi_R$), its subpartitioning is totally determined by cAMP, consistent with ref.[9,22,23], because, from (7), at steady-state

$$\frac{\phi_C}{\phi_A} = \frac{\beta_C([\text{cAMP}])}{\beta_A} \tag{S1.88}$$

The approximation of a constitutive A sector can be useful to explore the relationship between cAMP and ppGpp under anabolic limitation (A-LIM), extending the perspective shown in Fig. 2C. Under this type of growth limitation, the anabolic sector expands ($\phi_A$ increases), while the catabolic sector shrinks ($\phi_C$ decreases). The ribosomal sector reduces as well ($\phi_R$ decreases)[22,23,27]; therefore the P sector as a whole (A+C) must expand with increasing degree of limitation, because the sum $\phi_R + \phi_P$ must remain constant under the assumption of a constant Q sector. This expansion cannot be driven by cAMP activation, which would result in an increase in C at the expense of A. Therefore, repression of the R sector by increased ppGpp levels is necessary to indirectly upregulate metabolism. At the same time, cAMP levels must decrease to favor A over C, consistent with observations that cAMP levels decline under anabolic limitation[22,23]. This anticorrelation between ppGpp and cAMP is illustrated by the solid green arrow in Fig. 6. As explained in the main text, we predict the displacement from the C-LIM line in the cAMP-ppGpp plane will lead to a divergence from the C-LIM ribosome relationship (green dashed arrow in Fig. 6), similarly to what we observed under RelA overexpression.

While this simplified model captures large scale proteome allocation changes, we note that treating the A sector as purely constitutive can only be an approximation. Experimental data show that constitutive proteins are upregulated under catabolic limitation (e.g., Balakrishnan et al.[1]), whereas anabolic proteins are downregulated[16,22,23]. This discrepancy indicates that additional regulatory mechanisms govern anabolic genes expression, as already suggested by other works in the field[23,27].

#### 5.2 Translational inhibition

Another type of growth limitation is translational inhibition (R-LIM)[16]. A well-studied example is the antibiotic chloramphenicol (Cm)[9,24,28], which functions by stalling ribosomes[28]. Under Cm, the ribosomal fraction $\phi_R$ increases, and the elongation rate of the active (non-stalled) ribosomes $\epsilon$ also increases. Importantly, this increase follows the C-LIM relationship $\epsilon(\phi_R)$ (Fig. S4), as shown by Dai et al.[28]. Furthermore, Wu et al.[24] showed that Cm lowers ppGpp, and that this decrease continues to follow the wild-type relationship with elongation rate, $\epsilon(g)$ (Fig. 4A). Taken together, these two conserved relationships imply that Cm must also conserve the C-LIM mapping from ppGpp to ribosomal fraction $\phi_R(g)$:

$$g \underset{\text{Wu 2022}}{\overset{g(\epsilon)}{\longleftrightarrow}} \epsilon \underset{\text{Dai 2016}}{\overset{\epsilon(\phi_R)}{\longleftrightarrow}} \phi_R$$

This mapping is represented by the solid red arrow in Fig. 6.

In our model, $\phi_R$ is an function of the pair $(c, g)$, and it is decreasing in both variables (see equation (9)). Thus the C-LIM ppGpp-ribosome relationship $\phi_R(g)$ can be conserved if and only if the C-LIM cAMP-ppGpp relationship $c(g)$ is conserved as well. Based on the assumption that Cm preserves the C-LIM ribosome-ppGpp law, we predict it should also preserve the C-LIM cAMP-ppGpp one, as shown by the dashed red arrow in Fig. 6.

#### 5.3 Expression of useless proteins

Expressing useless proteins is commonly used to investigate resource allocation[9,22,27]. This perturbation can be incorporated into our model by introducing a sector corresponding to useless proteins, along with their mRNAs and genes, consistent with the framework of You et al.[22]. The ribosome allocation equation (7) can then be written as

$$\phi_R = \frac{\beta_R(g)}{\beta_R(g) + \beta_P(c) + \beta_U(x)} \phi_{max}, \tag{S1.89}$$

where $\beta_U$ is the coarse grained promoter strength associated with the useless gene, which is typically controlled by an inducer $x$ such as IPTG[9,22]. The parameter $\phi_{\text{max}}$ is identical to that in the wild type, since the housekeeping sector is assumed to be unaffected by the overexpression of useless proteins[9,22].

As evident from the above equation, varying the inducer concentration $x$ modulates the ribosomal fraction $\phi_R$. Although the impact of unneeded proteins on second messengers has not been quantified experimentally, it is important to note that, with increasing inducer, ribosome content could change even without any variation in cAMP or ppGpp (with constant $\beta_R(g)$ and $\beta_P(c)$). In this idealized scenario, ribosome content and ppGpp levels could become entirely uncorrelated, as indicated by the dashed black arrow in Fig. 6.
